# Leveraging the subgenus category to address genus over-splitting exemplified with *Prescottella* and other recently proposed *Mycobacteriales* genera

**DOI:** 10.1101/2025.04.27.650868

**Authors:** Jorge Val-Calvo, Mariela Scortti, José A. Vázquez-Boland

## Abstract

Three related circumstances are compromising the stability of prokaryotic taxonomy and nomenclature, with a significant impact in the field of pathogenic microorganisms: (i) the growing trend of subdividing existing genera by arbitrary phylogenomics-based demarcations, creating an increasing number of new genera; (ii) the priority of databases towards the last validly published names; and (iii) the irreversibility of new names in databases even if changes in taxonomic opinion support reverting to previous classifications. Given the understandable end-user reluctance to name changes affecting well-known pathogens, parallel nomenclatures coexist, creating confusion. Here, we address this problem by exploiting the subgenus category to form new combinations that can be adopted by databases as per their latest validly published name priority policy. The proposed approach consists in lowering to subgenus rank the new genera arising from genus splits considered to be unwarranted. According to the Prokaryotic Code, the species in question would be designated with their previous generic synonym and, in parentheses, the subgenus name (which would correspond to the latest synonym used by the databases). Being the subgenus an optional taxonomic category, it can be omitted, but its use may facilitate the mapping of the synonyms in databases and the literature. For illustration, we apply the strategy to recent genus splits in the *Mycobacteriales*, specifically the new nested genus *Prescottella* within the rhodococcal radiation and the several new genera into which *Mycobacterium* was subdivided.

---

A key underpinning principle in prokaryotic nomenclature is to aim at the stability of names (principle 1.1. of the International Code of Nomenclature of Prokaryotes [ICNP]) [1]. A stable nomenclature is essential for the effective study of microbiology, communication of microbiological knowledge, and traceability of microbial species in the literature. This notion is most evident in medicine, clinical microbiology and public health where microbial name changes, in addition to creating confusion, may lead to identification errors, misdiagnoses or inaccurate risk assessments potentially delaying or preventing appropriate treatment and control measures. However, taxonomic and nomenclatural stability is being compromised by the growing practice of subdividing existing genera into multiple new genera [2–10].

The genus over-splitting trend is a paradoxical consequence of the implementation of a genomics-based taxonomy and subjective application of genome similarity thresholds in the genus rank demarcation. This has been recently documented in a taxonomic study of the order *Mycobacteriales* [11] affected by two significant examples of genus fragmentation: the controversial [12–14] 2018 subdivision of *Mycobacterium* into five genera [6]; and the creation in 2022 of the nested genus *Prescottella* for the rhodococcal sublineage containing *Rhodococcus equi* [15]. The latter made the genus *Rhodococcus* paraphyletic, implying the need for creating additional genera for each of the other major rhodococcal sublineages of equal rank in order to ensure internal taxonomic coherence within the rhodococcal monophyletic radiation. Using a novel network analysis-aided approach for non-subjective taxonomic rank demarcation, the *Mycobacteriales* study showed that the aforementioned genus splits resulted from elevation of the intra-generic sublineages to genus rank by arbitrary application of genomic relatedness index (GRI, aka OGRI [16])-based demarcation boundaries [11]. Specifically, in the case of the proposed novel *Prescottella* genus, this involved shifting the genome-aggregate average amino acid identity (AAI) demarcation threshold to 74-75% [15], significantly deviating from the 65% threshold recognized as a robust genomic discreteness standard for genus definition in both natural isolates and metagenomic sequences [17–20]. There was also inconsistent use of other GRIs, such as values well above the proposed 50% for genus demarcation based on the percentage of conserved proteins (POCP) [21].

The arbitrary subdivision of established monophyletic genera is possible due to the absence of standardized genome-based demarcation guidelines and the freedom of taxonomic thought (Principle 1.4 of the ICNP [1]). Upon valid publication, the resulting new names are immediately relayed by all major nucleotide databases, notably the National Center for Biotechnology Information (NCBI)’s taxonomy browser and sequence repositories (GenBank). The information is then automatically mirrored by other main gene/genome databases, such as the European Nucleotide Archive (ENA) from the European Molecular Biology Laboratory (EMBL)-European Bioinformatics Institute (EBI), or the DNA Data Bank of Japan (DDBJ), under the backdrop of the International Nucleotide Sequence Database Collaboration (INSDC; http://www.insdc.org/) [22]. Indeed, one of the goals of the INSDC initiative is to use a unified taxonomy across all databases based on sequence information (https://ncbi.nlm.nih.gov/genbank/collab/) [23].

## DATABASE PRIORITY FOR LATEST VALIDILY PUBLISHED NAMES

Since according to ICNP rules both the new and earlier names are legitimate, they are synonyms [1] and thus the users are free to use the previous name (or any other validly published earlier synonym) [1, 24–26] if in disagreement with the taxonomic and nomenclatural changes (which strictly speaking only reflect a taxonomic opinion, by those proposing the changes). Additionally, the Taxonomy Project/NCBI Taxonomy database states that “they are not an authoritative source for nomenclature or classification” (https://www.ncbi.nlm.nih.gov/Taxonomy/Browser/wwwtax.cgi) [27, 28]; only the ICNP and the International Committee on Systematics of Prokaryotes (ICSP) have this prerogative. However, in practice the NCBI Taxonomy and mirror databases introduce a powerful “preference” bias toward specific names/synonyms, in two ways:

First, because of NCBI’s penetration and dominant position as a trusted public biosciences information repository, the taxonomic names they use are potentially perceived as representing the “officially-sanctioned” name for the species. This is likely to be so for most database users, who are not necessarily familiar with ICNP’s rules of synonymy. The NCBI itself explicitly recognizes the concept of “preferred name” in their Taxonomy Browser (https://www.ncbi.nlm.nih.gov/books/NBK53758) [29], favoring particular combinations vs other legitimate names (synonyms). For example, *Mycolicibacterium smegmatis* is indicated as the preferred name for *Mycobacterium smegmatis*, despite the mycobacterial nomenclature having been emended in 2021 by reclassifying the four additional genera created in 2018 back into a single genus *Mycobacterium* [14, 30, 31]. Similarly, the NCBI still considers *Prescottella equi* as the preferred name instead of *Rhodococcus equi* (see the entry in https://www.ncbi.nlm.nih.gov/Taxonomy/TaxIdentifier/tax_identifier.cgi), despite a recent change in taxonomic opinion having reclassified the species back into the genus *Rhodococcus* [11].

Second, the NCBI have adopted as a policy to use for their “primary name” (or “preferred name”, i.e. the name chosen out of all synonyms as the designated label for the TaxNode and its TaxID) [27], the latest validly published combination, despite the ICNP not stipulating that the latter should be treated as the correct name of a species vs all its synonyms. This is explained in Margos et al. [25] where they quote the following statement from NCBI: “Priority is given to taxonomic names validly published under the ICNP. In the case of two validly published names, one being a new combination of an earlier name, priority is given in the NCBI taxonomy database to the latest validly published name.” The application of this “priority to latest validly published name” policy by databases has two important practical consequences.

One is that it negates the possibility that changes in taxonomic opinion based on new evidence, better data or scientific advances –which are published as emendations and duly notified in “Lists of Changes in Taxonomic Opinion” [32]– are adequately reflected in the databases. This makes any proposed new combination virtually irreversible, thus in effect undermining the freedom of taxonomic thought or action. This NCBI policy has a major negative impact in situations where further taxonomic research determines that it would be more adequate to revert to an earlier classification. This applies in particular when subdivisions of well-established monophyletic genera which led to the creation of new generic names appear to be questionable, as the above discussed creation of the nested genus *Prescottella* within the *Rhodococcus* genus radiation or the *Mycobacterium* five-genus split [11, 14].

Secondly, if new species are described using the earlier genus synonym (in accord with current changes in taxonomic opinion that revert the genus splitting) instead of the latest validly published name used in the databases, NCBI Taxonomy considers their taxonomic check status as “inconclusive” (see e.g. https://www.ncbi.nlm.nih.gov/datasets/genome/GCF_963378085.1/, accessed 7 April 2025) or lists the species with the earlier genus name between square brackets with the caveat “awaits appropriate action by the research community to be transferred to another genus” (i.e. the one with the latest validly published name) (see e.g. https://www.ncbi.nlm.nih.gov/Taxonomy/Browser/wwwtax.cgi?mode=Info&id=3064284&lvl=3 &lin=f , accessed 7 April 2025). Thus, the NCBI appears to be demanding a publication that would formalize a novel combination that uses the latest validly published name for the genus even if against current taxonomic opinion. In other words, the “latest validly published name priority” policy the NCBI seems to be not only interfering with the freedom of taxonomic thought or action but also unduly influencing the development of phylotaxonomic research and microbial science.

## USE OF THE SUBGENUS TAXONOMIC RANK TO RECTIFY ARBITRARY GENUS SUBDIVISIONS

A possible strategy to circumvent the inconveniences derived from the databases’ practice of giving priority to the latest validly published names in cases where a subsequent change in taxonomic opinion corrects an arbitrary genus split would be to treat the taxon reversions as new combinations. This in our opinion is possible by taking advantage of the subgenus category, which, although seldom used, is recognized as a valid taxonomic rank by the ICNP [1].

Essentially, the approach consists in reclassifying to subgenus category any new genera resulting from monophyletic genus splits found to be taxonomically questionable. In this way, new taxons/combinations would be created which, as per ICNP’s subgenus notation [1], would carry both the earlier (pre-split) taxonomic name as generic designation and, in parenthesis, the latest proposed genus name that is being replaced, thus ensuring the straightforward identification and traceability of the different synonyms. This approach is illustrated below with the recently proposed nested rhodococcal genus *Prescottella* Sangal *et al*. 2018.

## DEMARCATION OF RHODOCOCCAL SUBGENERA

To accurately define the subgenus circumscriptions, we utilized our recently reported phylogenomic approach for taxon demarcation based on normalized tree clustering and network analysis of GRI and Maximum Likelihood (ML) distance (MLD) matrices. The approach was originally developed for genus delineation, but it was found to be applicable as a general taxonomic rank demarcation tool [11]. To minimise demarcation subjectivity, the method involves the uniform application of the same tree clustering/network graph partitioning thresholds across a sufficiently wide taxonomic context, not just the specific circumscription under study. This was the order level for genus delineation, using the classical/pre-genomic genera as demarcation reference to ensure taxonomic and nomenclatural continuity [11]. For *Rhodococcus* subgenus demarcation we are using here the entire *Nocardiaceae* radiation as taxonomic context, including the recently proposed *Hoyosellaceae* and *Tomitellaceae* families (which together form a major line of descent within the *Mycobacteriales* [11, 33]). The analyses included a total of nine genera: *Rhodococcus*, *Rhodococcoides*, *Nocardia*, *Antrihabitans*, *Tomitella*, *Hoyosella* and *Lolliginicoccus* (recently described within the family *Hoyosellaceae* [34]), plus the monospecies genera *Aldersonia* and *Skermania* (**Fig. S1**, **Suppl. dataset**).

We began by constructing a detailed ML phylogeny using a total of 175 genomes from all the species with available sequences within the study’s taxonomic context. To have a more comprehensive representation of the diversity within the *Rhodococcus*/*Rhodococcoides* radiation, the analyzed genomes also included 31 unclassified *Rhodococcus* spp. For the latter, one genome per each main terminal branching in the *Rhodococcus* spp. genomic Blast dendrogram available at the NCBI (https://www.ncbi.nlm.nih.gov/genome/?term=txid192944, accessed November 2022) was selected using as a filter an average nucleotide identity (ANI) ≥95% (the cutoff for species delineation [18, 35–37] (**Fig. S1**, **Suppl. dataset**). The TreeCluster programme “Max Clade” clustering method [38] was then used to partition the ML tree into discrete clusters based on evolutionary distances (branch lengths) and tree topology (phylogenetic relationships). To avoid clustering biases caused by branch differences in evolutionary rate, prior to the ML tree clustering, the branch lengths were normalised according to relative evolutionary distance (RED) using the PhyloRank package [39] (**Fig. 1**). For the taxonomic context under study, a TreeCluster threshold *t* = 0.95 recapitulated the genus structure of the *Nocardiaceae* radiation [11]. Decreasing the *t* value to 0.75 partitioned the tree into clusters which in the rhodococcal radiation roughly corresponded to the major intrageneric sublineages (**Fig. 1**). These analyses yielded the following conclussion:

i. Four major monophyletic lines of descent (designated by Arabic numerals in **Fig. 1**) can be identified within the genus *Rhodococcus* (2023 emendation [11, 40]) when taking as a partition reference the “Prescottella” sublineage (aka sublineage no. 2 or “equi” clade). For reasons of internal taxonomic consistency, all these sublineages should be considered as *Rhodococcus* subgenera if the *Prescottella* circumscription is classified under this taxonomic category.
ii. Two sublineages hierarchically equivalent to the above are also observed in the genus *Rhodococcoides* [11] and, accordingly, could also be treated as subgenera.
iii. Using the same genus and subgenus tree partitioning criteria, the novel genus *Lolliginicoccus* [34] does not warrant independent genus status (would not even qualify as a *Hoyosella* subgenus) (**Fig. 1**).
iv. The tree clustering analyses confirm the strong homogeneity of the *Nocardia* genus [11], with all its main sublineages radiating at short genetic distances well below the subgenus demarcation threshold (**Figs. 1 and S1**).

**Fig. 1.**
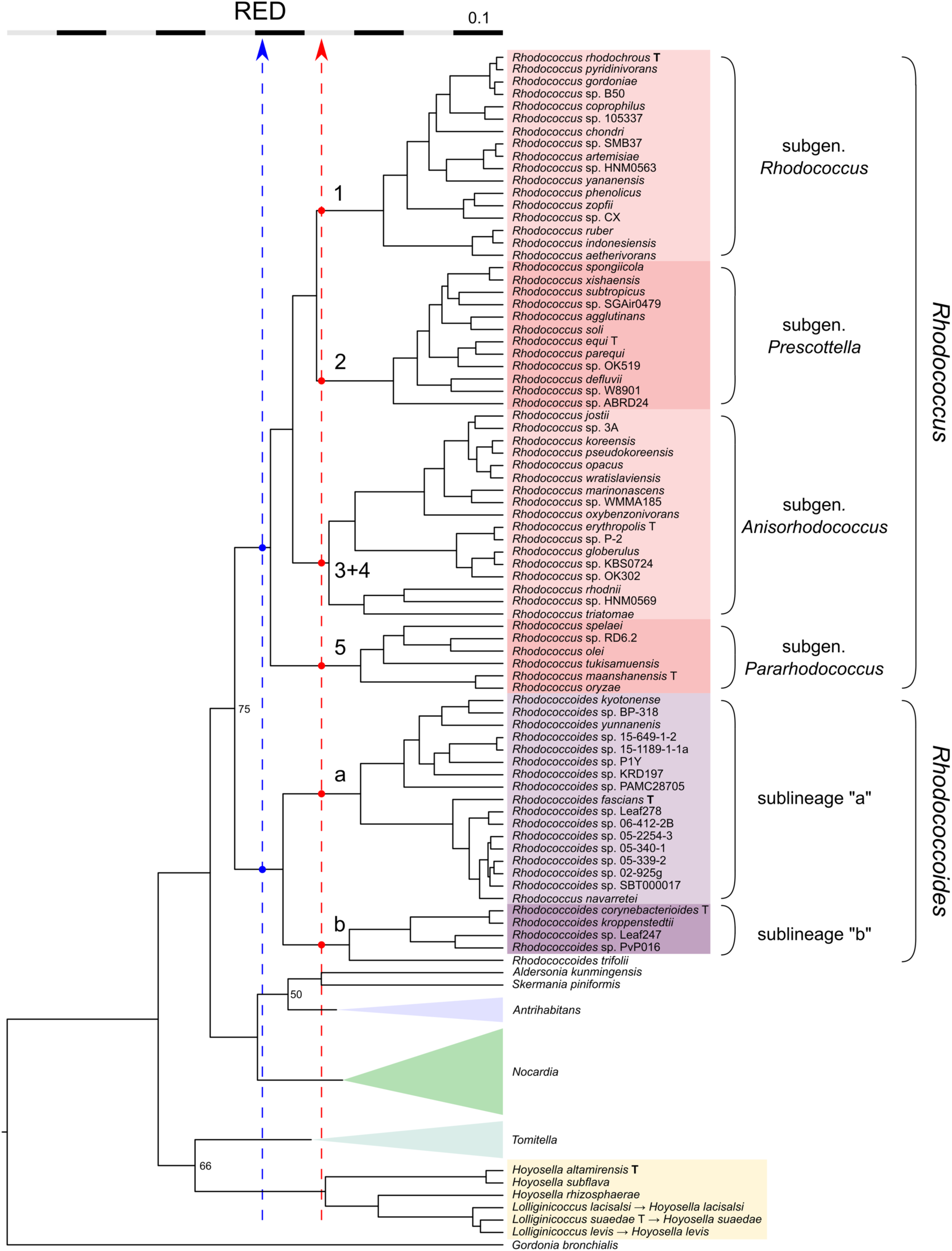
Relative evolutionary distance (RED)-normalized phylogenomic ML tree of the families *Nocardiaceae*, *Hoyosellaceae* and *Tomitellaceae*. See **Fig. S1** for the non-normalized version of the tree and details of the phylogenetic analysis. The genera *Nocardia*, *Antrihabitans* and *Tomitella* where no taxonomic changes are proposed are collapsed. The dashed lines indicate the TreeCluster partitioning thresholds for taxonomic context-uniform genus (blue) and subgenus (red) definition; the intersections with the rhodococcal radiation are indicated by dots. Branches of main sublineages within the genera *Rhodococcus* (shaded red) and *Rhodococcoides* (shaded pink) are indicated by numbers [45] and lower-case letters, respectively. On the right, the circumscriptions of the proposed subgenera are indicated by braces. Note in the tree (i) that the genus-level partition does not support the proposed new genus *Lolliginicoccus* within the *Hoyosella* radiation; and (ii) the recently described species *Rhodococcus navarretei* is located within sublineage “a” of the *Rhodococcoides* genus radiation. Bootstrap values below 75 (1,000 replicates) are shown. Tree plotted using FigTree v1.4.4 (http://tree.bio.ed.ac.uk/software/figtree/).

## TAXONOMIC NETWORK ANALYSIS

Next, the tree clustering-based subgenus rank demarcations were validated using network analysis of MLD and GRI matrices [11]. The MLD matrix was constructed with the **Figs. 1** (and **S1**) phylogenetic tree dataset. The GRI matrices used were based on AAI [17, 19, 20] and aligned fraction of orthologous genes (AF) scores [41, 42]. In these analyses, the MLD/GRI pairwise comparison matrices are used to generate a correlation matrix which is then three-dimensionally (3D) represented in a network graph to visualize the taxonomic relationships [11]. The network graphs were generated with the open-source software Graphia, an improved version of Biolayout used in our previous *Mycobacteriales* study with better compatibility, correlation analysis of high dimensional numerical matrices and graphic environment [43, 44]. To explore the degree of relatedness of the species included in the analysis, network graphs with increasing fragmentation were generated along a correlation/clustering threshold (ct) range.

As observed in our previous *Mycobacteriales* study [11], a specific range of MLD/GRI ct cutoffs recapitulated the genus structure of the taxonomic context under analysis, isolating the genera as discrete subnetworks (**Fig. 2**, left). An exception was the *Lolliginicoccus* circumscription, which was completely embedded in the *Hoyosella* genus subnetwork in all three genus-level MLD/GRI network graphs. This result was consistent with the tree clustering data (**Fig. 1**) and AAI scores with the *Hoyosella* genomes exceeding the ≤65% standard for genus definition [17–20] (mean value, 69.26%), further supporting that the *Lolliginicoccus* genus taxon is not justified.

**Fig. 2.**
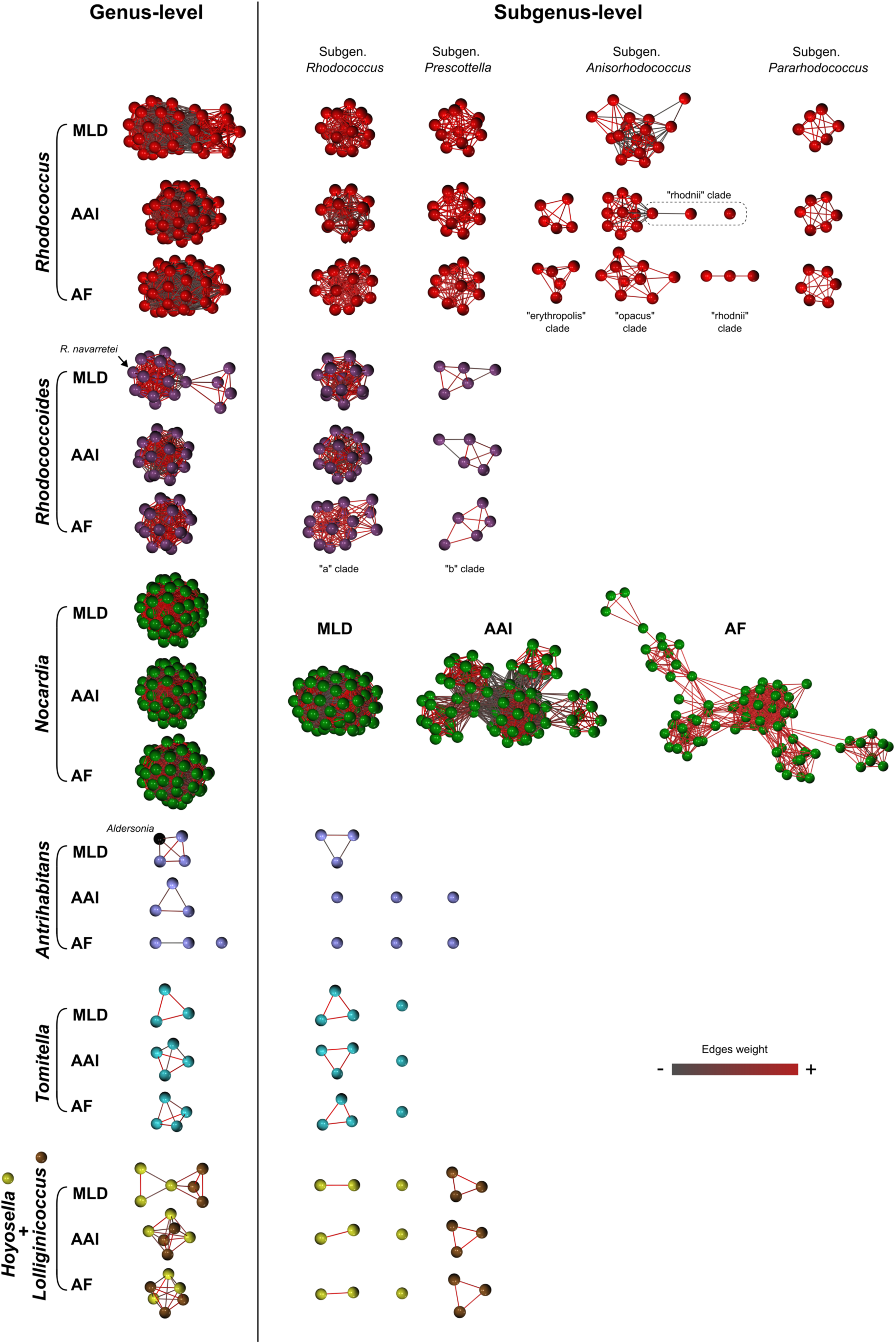
Phylogenomic 3D network analysis based on ML distance (MLD) and GRI (AAI, AF) matrices using **Fig. 1** dataset. Each node represents a genome and interconnecting edges show relatedness between nodes above the ct correlation/clustering threshold (similarity weight shown as grey [weaker] to red [stronger] gradient). The stronger the genomes/species are related, the closer their nodes sit in the network and the stabler their connectivity remains upon increasing the ct. Left panels, genus level partitions; ct values: MLD 0.670, AAI 0.410, AF 0.525. Right panels, subgenus level partitions; ct values: MLD 0.825, AAI 0.720, AF 0.810. The ct thresholds are set according to the taxonomic context under analysis; the specific ct values for genus- or subgenus-level partitions vary depending on the genome datased used. Note in the genus-level graphs that *Rhodococcus navarretei* Carrasco *et al*. 2024 [58] is embedded in the *Rhodococcoides* network, indicating it is part of this genus circumscription (see also **Fig. 1** legend). Note also that the monotypic genus *Aldersonia* remains connected to the *Antrihabitans* subnetwork in the MLD-based genus graphs. According to previous taxonomic network analyses, *Aldersonia kunmingensis* DSM 45001^T^ and *Skermania piniformis* DSM 43998^T^ (also a monotypic genus) occupy an intermediate, linking position between the *Rhodococcus* and *Nocardia* clusters, with *Aldersonia* connecting with both *Antrihabitans* and *Skermania* [11]. This is also shown in **Fig. S2** where a lower ct was applied to **Fig. 2** dataset, enabling the visualisation of supragenus-level relationships.

By increasing the ct cutoffs to isolate the *Prescottella* sublineage circumscription in a self-contained subnetwork, additional discrete subnetworks (putative subgenera) are formed which overall correspond to each of the previously identified rhodococcal sublineages (**Fig. 2**, right). The same occurs with the two identified main *Rhodococcoides* sublineages (ref. [11] and herein), whereas all the species of the genus *Nocardia* remain connected in a same subnetwork (**Fig. 2**, right), closely mirroring the ML tree clustering partitions (**Fig. 1**).

Only in one instance, i.e. the AAI/AF-based network graphs of *Rhodococcus* sublineage no. 3+4, the subgenus-level partitions distributed the corresponding circumscription into more than one subnetwork (**Fig. 2**). The 3+4 “subgeneric” subnetworks correspond to the main internal branches of this sublineage (see **Fig. 1 and S1**), which differ in genome size and topology (“erythropolis” clade, circular genomes of ≈6 Mbp; “opacus/jostii” clade, large linear genomes of >8-9 Mpb; “rhodnii” clade, smaller circular genomes of 4-5 Mbp). This internal genomic heterogeneity is unique among the rhodococcal lines of descent and may account for the observed separation of the AAI/AF-based subgenus-level network graphs of the 3+4 sublineage.

## PROPOSAL TO LOWER THE GENUS *PRESCOTTELLA* SANGAL *ET AL*. 2022 TO SUBGENUS RANK

To address the problematic co-existence of two different names for the same pathogen −*Rhodococcus equi* and the later synonym *Prescottella equi*−, based on our phylogenomic analyses we propose to reclassify the genus *Prescottella* Sangal *et al.* 2022 [15] as a subgenus of the genus *Rhodococcus* Zopf 1891 (Approved Lists 1980) emend. Val-Calvo and Vazquez-Boland 2023 [40]. For the sake of internal taxonomic consistency, we propose to elevate in parallel the other main rhodococcal monophyletic sublineages, as identified in this work and previous phylogenomic studies [11, 45], to subgenera of the genus *Rhodococcus*. All *Rhodococcus* subtaxons defined in this study have average inter-subgeneric AAI values comprised between the 65% standard for genus definition and 73% (**Fig. 3**).

**Fig. 3.**
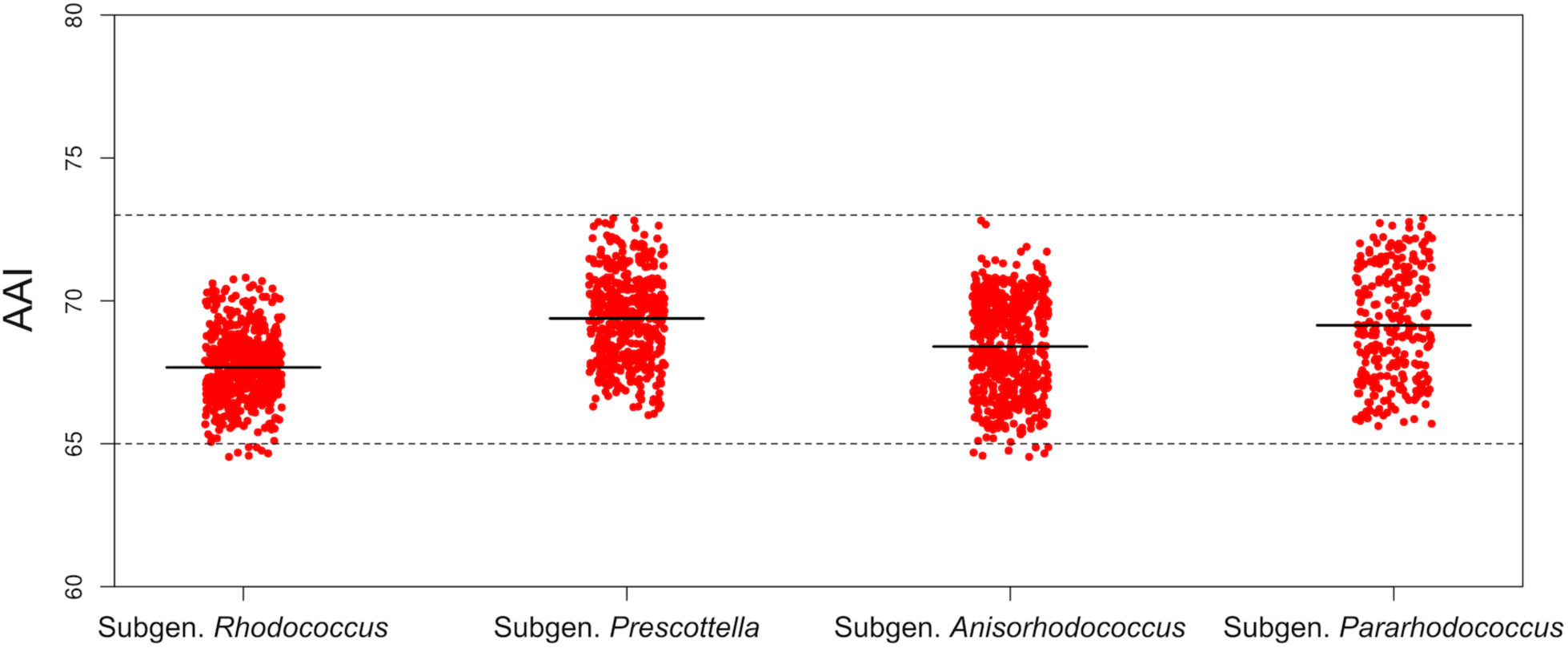
Scatter plots of *Rhodococcus* inter-subgenus AAI scores. The horizontal dashed line indicates the standard 65% AAI genus demarcation standard [17, 18, 20]. The rhodococcal subgenus demarcation boundary is AAI = 73%. The average AAI value for each subgenus is represented by a black horizontal line.

Our approach to subgenus definition is conservative, i.e. we chose a taxon demarcation threshold within the *Prescottella*-level range which minimised the creation of new names. Accordingly, the *Rhodococcus* monophyletic sublineage 3+4, which contains three lines of descent within the *Prescottella*-level subgenus definition range (i.e. the “erythropolis”, “opacus/jostii” and “rhodnii” clades) (**Fig. 1 and S1**), was considered as a single subgenus. This taxon contains species of different genome size and topology (see above) and hence we propose for it the subgenus name “*Anisorhodococcus*”. Species with different genome size being classified within a same genus or, as in this case, subgenus, is perfectly possible. While linked to bacterial phylogeny at a broader scale [46], genome size (which influences genome topology, i.e. larger actinomycetal genomes tend to be linear [47]) may vary within a specific group of closely related bacteria due to niche-adaptive gene content expansion or contraction and, thus, does not have strict taxonomic value [48–50].

The *Rhodococcus* subgenus nomenclature was proposed in accord with the precepts of the ICNP [1], as follows. Rule 49 states that “when a genus is lowered in rank to subgenus, the original name must be retained unless it is rejected under the Rules”. Rule 39a states that “If a genus is divided into two or more genera or subgenera, the generic name must be retained for one of these”. Finally, Rule 39b states “When a particular species has been designated as the type, the generic name must be retained for the genus which includes that species.” Thus, the name *Prescottella* is used here to designate the subgenus that corresponds to sublineage 2 containing *R. equi*, and *Rhodococcus* is retained as the name of sublineage 1 containing the type species of the genus *Rhodococcus*, *Rhodococcus rhodochrous*. To minimise subjectivity in the rank demarcations, we used the tree clustering and MLD/GRI-based network analysis approach which we had previously applied to reassess the *Mycobacteriales* taxonomy [11]. Using these criteria, the two sublineages within the genus *Rhodococcoides* Val-Calvo and Vázquez-Boland 2023 would also warrant subgenus rank. However, we are not proposing for these the creation of new subgenera; we restrict ourselves to the genus *Rhodococcus* “*sensu stricto*” (i.e. as recently emended by Val-Calvo and Vazquez-Boland) [11, 40], as the main purpose of this study was to address the confusion (and taxonomic implications) created by the proposal of the nested genus *Prescottella* within the rhodococcal monophyletic radiation.

## APPLICATION TO THE *MYCOBACTERIUM* FIVE-GENUS SPLIT

Another obvious candidate for application of the subgenus strategy delineated here is the subdivision of the original monophyletic genus *Mycobacterium* into five different genera proposed by Gupta et al. in 2018 [6]. The new mycobacterial genus names that resulted from this taxonomic change are used by databases but largely rejected by the end-user community [12–14, 31].

As a first step, we assessed the taxonomic consistency of Gupta et al. [6] *Mycobacterium* subdivision by mapping the five new genera on an ML tree constructed using 160 mycobacterial genomes (**Fig. S3, left**). The tree shows that, apart from an early diverging clade (sublineage “v”, assigned to the genus *Mycobacteroides*), the bulk of the *Mycobacterium* (*sensu lato*) circumscription splits into two major branches. One of them was subdivided by Gupta into three unequal genus partitions: one corresponded to the larger of the two sister clades that make up this branch (labelled sublineage “i” in the tree), contained the type species, *Mycobacterium tuberculosis*, and was assigned to an emended genus *Mybobacterium* [6]; the sister clade was further subdivided by Gupta et al. [6] into two novel genera, *Mycolicibacter* (sublineage “iii”) and *Mycolicibacillus* (sublineage “iv”). The second major branch (labelled sublineage “ii” in oour tree) was assigned by Gupta et al. [6] to the novel genus *Mycolicibacterium* (**Fig. S3, left**). From this analysis it is evident that the genus partitions proposed by Gupta et al. [6] are not homogeneous in terms of rank or position in the mycobacterial tree hierarchy.

The lack of consistency of Gupta *et al.* mycobacterial five-genus split is most evident in a RED-normalised ML tree (**Fig. S3, right**). Here, application of a context-homogeneous partitioning [11] that preserves the more distal *Mycolicibacter* and *Mycolicibacillus* demarcations would subdivide the mycobacterial radiation into a total of 14 taxa (subgenera) of an equivalent rank (**Fig. S3, right**). On the other hand, if we take *Prescottella* as the subgenus partitioning reference across the broader *Mycobacteriales* taxonomic context [11] (**Fig. S4),** *Mycobacterium* (*sensu lato*) would be subdivided into three subgenera (labelled A, B, C) that would correspond to Gupta *et al*. (A) *Mycobacterium* (emend.) + *Mycolicibacter* + *Mycolicibacillus*, (B) *Mycolicibacterium*, and (C) *Mycobacteroides*. Given the impossibility of reconstituting Gupta et al. mycobacterial five-genus split using a context-uniform demarcation approach, there is no point in applying our taxon delineation methodology in this case. We therefore propose to approach the problem in a pragmatic manner, by simply lowering the new mycobacterial genera proposed by Gupta *et al*. [6] to subgenus rank. In this way, the entire mycobacterial radiation (with the exception of the *Mycobacteroides* circumscription, see below) would again be designated with the same generic name *Mycobacterium*, while each of the subgenera contained within it would retain the (generic) names coined by Gupta et al. [6] currently in use in the databases −thus allowing users to quickly identify at a simple glance the relationship between the two nomenclatures.

The *Mycobacteroides* clade (mycobacterial sublineage “v”; **Fig. S3**) requires special consideration. Our recent taxonomic analysis of the *Mycobacteriales* found that, in contrast to the other mycobacterial genera proposed by Gupta *et al*. [6], the early diverging mycobacterial clade “v” containing *Mycobacteroides abscessus* ATCC 19977^T^ would warrant independent genus status [11]. The study implemented a taxonomic-context uniform demarcation strategy across the entire *Mycobacteriales* circumscription and used the classical (pre-genomic) genera as taxon partitioning reference to ensure continuity and consistency in taxonomy and nomenclature. Using this approach, the *Mycobacteroides* clade appeared as a robust genus by all partitioning methods used, entirely equivalent in rank and hierarchical position to other established *Mycobacteriales* genera such as *Gordonia*, *Williamsia*, *Tsukamurella* or *Hoyosella* (some of which are also early diverging branches of monophyletic radiations forming other genera, as for example *Williamsia* or *Tsukamurella* in relation to *Gordonia*) [11]. The separation of *Mycobacteroide*s as a separate genus is also supported by an AAI score ≤65% with the other mycobacterial clades and *Mycobacteriales* genera (whereas the cutoff for taxon discreteness for the four proposed *Mycobacterium* subgenera is ≤73%, the same as that for the rhodococcal subgenera, see above) (**Fig. S5**). The possible separation of *Mycobacteroides* as an independent mycobacterial genus was also suggested in the recent taxonomic study by Meehan *et al*. that proposed the reconstitution of the genus *Mycobacterium* [14]. Despite all this evidence, the mycobacterial community appears to remain unfavourable to an independent *Mycobacteroides* genus and prefers to consider its circumscription to be part of a single genus *Mycobacterium* encompassing the entire mycobacterial radiation [13]. For the sake of taxonomic consistency in genus demarcation across the *Mycobacteriales*, we are not proposing here to re-classify *Mycobacteroides* Gupta *et al*. 2018 emend. Val-Calvo and Vazquez-Boland 2023 as a subgenus of the genus *Mycobacterium*; we leave this to the consideration of the mycobacterial research community.

## TAXONOMIC CONCLUSIONS

The strategy of lowering the rank of the new genera created by arbitrary subdivision of a monophyletic genus to subgenus category offers, in our opinion, a reasonable “compromise” solution to the conundrum posed by the “latest validly published name priority” database policy. This practice potentially fixes *ad perpetuum* in the databases any latest validly published generic names unless replaced by new validly published combinations, in this case new subgeneric taxons as proposed here. Although the use of the subgenus rank was discouraged by the Judicial Commission of the ICSP [51], it remains a valid possibility in the Prokaryotic Code , and its application is essentially a matter of freedom of taxonomic thought, protected by the ICNP [1]. We believe that, for the purpose delineated here and if it remains restricted to this particular application, the use of the subgenus rank is acceptable, justified and useful. Taking this action would help mitigating any potential confusion generated by the recent genera subdivisions and reclassification of many prokaryotic species with well-known names under new genus designations, while also achieving the purpose of the genus rank, i.e. unifying under a same generic name a group of closely related microorganisms sharing the same recent ancestry and key biological attributes. Since *Rhodococcus* is a quite diversified genus, the availability of distinct subgenus names will also allow the easy distinction of the main rhodococcal sublineages. Currently, they designated by numbers [11, 45, 52, 53] or by the epithet/name of the prototype species [54–56], clearly highlighting the need for a specific nomenclature to refer to them in scientific studies −a need that we believe a subgeneric name would adequately satisfy. The very same rationale is also applicable to the mycobacterial sublineages (subgenera). It could be argued that proposing new subgenera may go against Principle 1.3 of the Prokaryotic Code “avoid the useless creation of names” [1]. However, since the subgenus is an optional taxonomic category (Rule 5b of the Code) [1], we anticipate that, in practice, the proposed subgeneric names will be rarely used and the bacteria classified under them will be mostly known and designated by their generic name, i.e. *Rhodococcus* and *Mycobacterium*.

Other taxonomic conclusions from this study include two taxon reclassifications. One affects the recently proposed genus *Lolliginicoccus* Miyanishi *et al.* 2023 [34], which we find is unjustified and in fact represents another example of the recent genus over-splitting trend. Consequently, of the current three *Lolliginicoccus* species, two need to revert to their basonyms/homotypic synonyms *Hoyosella lacisalsi* Yang et al. 2021 and *Hoyosella suaedae* Liu et al. 2021; while the third species, *Lolliginicoccus levis* Miyanishi *et al*. 2023, requires description as a new *Hoyosella* combination. The other reclassification concerns the recently described species *Rhodococcus navarretei*, which our ML phylogeny and taxonomic network analyses unambiguously place within the *Rhodococcoides* radiation. We therefore propose for the latter its reclassification as *Rhodococcoides navarretei* comb. nov.

## TAXONOMIC DESCRIPTIONS

### *Prescottella* (Sangal *et al*. 2022) subgen. nov

(Pres.cot.tel’la. N.L. fem. dim. n. *Prescottella*, in honor of John Prescott for his pioneering research contributions into the pathogenicity and epidemiology of *Rhodococcus equi*.) The description of this taxon is as given by Sangal *et al*. [15]. Previously described as a genus, it is lowered to a subgenus of the genus *Rhodococcus* Zopf 1891 (Approved Lists 1980) emend. Val-Calvo and Vazquez-Boland 2023. Core-genome phylogenies as well as RED-normalized tree clustering and GRI- and maximum likelihood distance-based network analyses all show that the Rhodococcus (subgen. *Prescottella*) species belong to an internal lineage of the genus Rhodococcus with intra-subgenus average AAI score of 83.6% and inter-subgenus AAI scores of ≤73%. It contains the species *Rhodococcus* (subgen. *Prescottella*) *agglutinans*, *defluvii*, *equi, parequi, soli, spongiicola, subtropicus,* and *xishaensis.* See **Table 1** for description of the new combinations.

**Table 1.** Descriptions of new *Rhodococcus* combinations.

| New name combinations | Basonym/Synonyms | Description and type strain |
| --- | --- | --- |
| ***Rhodococcus* (subgen. *Prescottella* Sangal *et al.* 2022) *agglutinans* comb. nov.**<br>(ag.glu'ti.nans. L. part. adj. *agglutinans*, sticking together, agglutinating) | **Basonym:** *Rhodococcus agglutinans* Guo *et al.* 2015<br>**Synonym:** *Prescottella agglutinans* (Guo *et al.* 2015) Sangal *et al.* 2022 | The description of this taxon is as given by Guo *et al.* [59]. The type strain has a G+C content of 69.2% and a genome (assembly) of $\approx 5.4$ Mbp.<br>**Type strain:** CFH S0262<sup>T</sup> (= CCTCC AB2014297<sup>T</sup> = KCTC 39118<sup>T</sup>). |
| ***Rhodococcus* (subgen. *Prescottella* Sangal *et al.* 2022) *defluvii* comb. nov.**<br>(de.flu'vi.i. L. gen. neut. n. *defluvii*, of sewage) | **Basonym:** *Rhodococcus defluvii* Kämpfer *et al.* 2014.<br>**Synonym:** *Prescottella defluvii* (Kämpfer *et al.* 2014) Sangal *et al.* 2022. | The description of this taxon is as given by Kämpfer *et al.* [60]. The type strain has a G+C content of 68.7% and a genome (assembly) of $\approx 5.13$ Mbp.<br>**Type strain:** Ca11<sup>T</sup> (= DSM 45893<sup>T</sup> = LMG 27563<sup>T</sup>). |
| ***Rhodococcus* (subgen. *Prescottella* Sangal *et al.* 2022) *equi* comb. nov.**<br>(e'qui. L. gen. masc. n. *equi*, of the horse) | **Basonym:** *Corynebacterium equi* Magnusson 1923 (Approved Lists 1980)<br>**Synonyms:** *Nocardia restricta* (Turfitt 1944) McClung 1974 (Approved Lists 1980); *Prescottella equi* (Magnusson 1923) Sangal *et al.* 2022 | The description of this taxon is as given by Goodfellow and Alderson [57] and Jones and Goodfellow [54]. The type strain has a G+C content of 68.8% and a circular genome (assembly) of $\approx 5.2$ Mbp.<br>**Type strain:** C7<sup>T</sup> (= ATCC 25729<sup>T</sup> = ATCC 6939<sup>T</sup> = CCUG 892<sup>T</sup> = CIP 54.72<sup>T</sup> = DSM 20307<sup>T</sup> = HAMBI 2061<sup>T</sup> = JCM 1311<sup>T</sup> = JCM 3209<sup>T</sup> = LMG 18452<sup>T</sup> = NBRC 101255<sup>T</sup> = NCTC 1621<sup>T</sup> = NRRL B- 16538<sup>T</sup> = VKM Ac-953<sup>T</sup>). |
| ***Rhodococcus* (subgen. *Prescottella* Sangal *et al.* 2022) *parequi* comb. nov.**<br>(par.e'qui. Gr. prep. *para*, beside, alongside of, near, like; L. gen. masc. n. *equi*, of a horse, specific epithet; N.L. gen. masc. n. *parequi*, resembling *Rhodococcus equi*) | **Basonym:** *Rhodococcus parequi* Vazquez-Boland *et al.* 2025 | The description of this taxon is as given by Vazquez-Boland *et al.* [45]. The type strain has a G+C content of 69% and circular genome with a size of $\approx 5.3$ Mbp.<br>**Type strain:** PAM 2766<sup>T</sup> (= CETC 30995<sup>T</sup> = NCTC 14987<sup>T</sup>). |
| ***Rhodococcus* (subgen. *Prescottella* Sangal *et al.* 2022) *solii* comb. nov.**<br>(so'li. L. gen. neut. n. *solii*, of/from the soil) | **Basonym:** *Prescottella solii* (Li *et al.* 2015) Sangal *et al.* 2022<br>**Synonym:** *Rhodococcus solii* Kämpfer *et al.* 2014 | The description of this taxon is as given by Li *et al.* [61]. The type strain has a G+C content of 68.5% and circular genome with a size of $\approx 5.5$ Mbp.<br>**Type strain:** DSD51W<sup>T</sup> (= KCTC 29259<sup>T</sup> = JCM 19627<sup>T</sup> = DSM 46662<sup>T</sup> = KACC 17838<sup>T</sup>). |
| ***Rhodococcus* (subgen. *Prescottella* Sangal *et al.* 2022) *subtropicus* comb. nov.**<br>(sub.tro'pi.cus. N.L. masc. adj. *subtropicus*, subtropical, implying that the type strain was isolated from subtropical zones) | **Basonym:** *Rhodococcus subtropicus* Lee *et al.* 2019<br>**Synonym:** *Prescottella subtropica* (Lee *et al.* 2019) Sangal *et al.* 2022 | The description of this taxon is as given by Lee *et al.* [62]. The type strain has a G+C content of 69.1% and a genome size (assembly) of $\approx 4.4$ Mbp.<br>**Type strain:** C9-28<sup>T</sup> (= DSM 107559<sup>T</sup> = KACC 19823<sup>T</sup>). |
| ***Rhodococcus* (subgen. *Prescottella* Sangal *et al.* 2022) *spongiicola* comb. nov.** | **Basonym:** *Rhodococcus spongiicola* Zhang *et al.* 2021 | The description of this taxon is as given by Zang *et al.* [63]. The type strain has a G+C content of 66.5% and circular genome with a size of $\approx 4.0$ Mbp. |
| (spon. gi.i'co. la. L. fem. n. spongia a sponge; L. suff. -cola (from L. masc. or fem. n. incola) inhabitant; N.L. masc. n. spongiicola inhabitant of sponge) |  | **Type strain:** LHW50502<sup>T</sup> (=DSM 106291<sup>T</sup> = CCTCC AA 2018033<sup>T</sup>). |
| ***Rhodococcus* (subgen. *Prescottella* Sangal *et al.* 2022) *xishaensis* comb. nov.**<br>(xi.sha.en'sis. N.L. masc./fem. adj. *xishaensis*, from Xisha Island, PR China, where the type strain was isolated) | **Basonym:** *Rhodococcus xishaensis* Zhang *et al.* 2021 | The description of this taxon is as given by Zhang *et al.* [63]. The type strain has a G+C content of 66.5% and circular genome with a size of ≈3.7 Mbp.<br>**Type strain:** LHW51113<sup>T</sup> (= DSM 106204<sup>T</sup> = CCTCC AA 2018034<sup>T</sup>). |
| ***Rhodococcus* (subgen. *Anisorhodococcus*) *erythropolis* comb. nov.**<br>(e.ry.thro.po.lis Gr. masc. Adj. *erythros*, red; Gr. fem. n. *polis*, a city; N.L. fem. n. *erythropolis*, red community) | **Basonym:** *Rhodococcus erythropolis* (Gray and Thornton 1928) Goodfellow and Alderson 1979 (Approved Lists 1980)<br>**Synonyms:** "*Mycobacterium erythropolis*" Gray and Thornton 1928; *Arthrobacter picolinophilus* Tate and Ensign 1974 (Approved Lists 1980); *Rhodococcus baikonurensis* Li *et al.* 2004; *Rhodococcus qingshengii* Xu *et al.* 2007; *Rhodococcus jialingiae* Wang *et al.* 2010; *Rhodococcus enclensis* Dastager *et al.* 2014; *Rhodococcus degradans* Švec *et al.* 2015 | The description of this taxon is as given by Goodfellow and Alderson [57] and Jones and Goodfellow [54]. The type strain has a G+C content of 62.5 % and a genome size of ≈6.6 Mbp.<br>**Type strain:** N11<sup>T</sup> (= ATCC 4277<sup>T</sup> = CIP 104179<sup>T</sup> = DSM 43066<sup>T</sup> = HAMBI 1953<sup>T</sup> = IEGM 7<sup>T</sup> = IFO 15567<sup>T</sup> = JCM 20419<sup>T</sup> = JCM 3201<sup>T</sup> = LMG 5359<sup>T</sup> = NBRC 15567<sup>T</sup> = NCIB 9158<sup>T</sup> = NCIMB 9158<sup>T</sup> = NCTC 13021<sup>T</sup> = NRRL B-16025<sup>T</sup> = VKM Ac-858<sup>T</sup>). |
| ***Rhodococcus* (subgen. *Anisorhodococcus*) *globerulus* comb. nov.**<br>(glo.be.ru.lus. N.L. masc. adj. *globerulus*, globular) | **Basonym:** *Rhodococcus globerulus* Goodfellow *et al.* 1985<br>**Synonym:** *Nocardia globerula* (Gray 1928) Waksman and Henrici 1948 (Approved Lists 1980) | The description of this taxon is as given by Goodfellow *et al.* [64] and Jones and Goodfellow [54]. The type strain has a G+C content of 61.7 % and a genome (assembly) size of ≈6.7 Mbp.<br>**Type strain:** R58<sup>T</sup> (= ATCC 25714<sup>T</sup> = CIP 104174<sup>T</sup> = DSM 43954<sup>T</sup> = IEGM 591<sup>T</sup> = IFO 14531<sup>T</sup> = JCM 7472<sup>T</sup> = NBRC 14531<sup>T</sup> = NRRL B-16938<sup>T</sup>). |
| ***Rhodococcus* (subgen. *Anisorhodococcus*) *jostii* comb. nov.**<br>(jost'i.i. N.L. gen. masc. n. *jostii*, of Jošt, in honor of Margrave Jošt Lucemburský (1351-1411), an important Czech ruler, from whose skeletal remains the strain was isolated) | **Basonym:** *Rhodococcus jostii* Takeuchi *et al.* 2002 | The description of this taxon is as given by Takeuchi *et al.* [65]. The type strain has a G+C content of 67 % and a linear genome with a size of ≈10 Mbp.<br>**Type strain:** IFO 16295<sup>T</sup> (= CCM 4760<sup>T</sup> = DSM 44719<sup>T</sup> = JCM 11615<sup>T</sup> = NBRC 16295<sup>T</sup>). |
| ***Rhodococcus* (subgen. *Anisorhodococcus*) *koreensis* comb. nov.**<br>(ko.re.en'sis. N.L. masc./fem. adj. *koreensis*, of or belonging to Korea, the country where the strain was isolated) | **Basonym:** *Rhodococcus koreensis* Yoon *et al.* 2000 | The description of this taxon is as given by Yoon *et al.* [66]. The type strain has a G+C content of 67.5 % and a linear genome (assembly) with a size of ≈10.3 Mbp.<br>**Type strain:** DNP 505<sup>T</sup> (= CIP 106721<sup>T</sup> = DSM 44498<sup>T</sup> = JCM 10743<sup>T</sup> = KCTC 0569BP<sup>T</sup> = NBRC 100607<sup>T</sup>). |
| ***Rhodococcus* (subgen. *Anisorhodococcus*) *marinonascens* comb. nov.** | **Basonym:** *Rhodococcus marinonascens* Helmke and Weyland 1984 | The description of this taxon is as given by Helmke and Weyland [67] and Jones and Goodfellow [54]. The type strain has a G+C content of 64.5 % and genome (assembly) size of ≈4.9 Mbp. |
| (ma.ri.no.na.scens. L. masc. adj. *marinus*, of the sea; L. pres. part. *nascens*, born; N.L. part. adj. *marinonascens*, born of the sea) |  | **Type strain:** 3438W<sup>T</sup> (= ATCC 35653<sup>T</sup> = CIP 104177<sup>T</sup> = DSM 43752<sup>T</sup> = IFO 14363<sup>T</sup> = JCM 6241<sup>T</sup> = NBRC 14363<sup>T</sup> = NRRL B-16940<sup>T</sup> = VKM Ac-1182<sup>T</sup>). |
| ***Rhodococcus* (subgen. *Anisorhodococcus*) *opacus* comb. nov.**<br>(o.pa.cus. L. masc. adj. *opacus*, shady, obscure, nontransparent) | **Basonym:** *Rhodococcus opacus* Klatte *et al.* 1995<br>**Synonyms:** *Rhodococcus percolatus* Briglia *et al.* 1996; *Rhodococcus imtechensis* Ghosh *et al.* 2006 | The description of this taxon is as given by Klatte *et al.* [68]. The type strain has a G+C content of 67.5 % and a linear genome with a size of ≈8.5 Mbp.<br>**Type strain:** DSM 43205<sup>T</sup> (= ATCC 51881<sup>T</sup> = CIP 104549<sup>T</sup> = IFO 16217<sup>T</sup> = JCM 9703<sup>T</sup> = NBRC 100624<sup>T</sup> = NBRC 16217<sup>T</sup>). |
| ***Rhodococcus* (subgen. *Anisorhodococcus*) *oxybenzonivorans* comb. nov.**<br>(xy.ben.zo.ni.vorans. N.L. neut. n. *oxybenzonum*, oxybenzone; L. pres. part. *vorans*, devouring; N.L. part. adj. *oxybenzonivorans*, oxybenzone-consuming) | **Basonym:** *Rhodococcus oxybenzonivorans* Baek *et al.* 2022 | The description of this taxon is as given by Baek *et al.* [69]. The type strain has a G+C content of 65.5 % and circular genome with a size of ≈8.0 Mbp.<br>**Type strain:** S2-17<sup>T</sup> (= JCM 32046<sup>T</sup> = KACC 19281<sup>T</sup>). |
| ***Rhodococcus* (subgen. *Anisorhodococcus*) *pseudokoreensis* comb. nov.**<br>(pseu.do.ko.re.en'sis. Gr. masc./fem. adj. I, false; N.L. masc./fem. adj. *koreensis*, ko.re.en'sis, of or belonging to Korea; N.L. masc./fem. adj. *pseudokoreensis*, false *R. koreensis*) | **Basonym:** *Rhodococcus pseudokoreensis* Kämpfer *et al.* 2023 | The description of this taxon is as given by Kämpfer *et al.* [70]. The type strain has a G+C content of 67.5 % and a linear genome (assembly) with a size of ≈9.9 Mbp.<br>**Type strain:** R79<sup>T</sup> (= CCM 9183<sup>T</sup> = DSM 113102<sup>T</sup> = LMG 32444<sup>T</sup>). |
| ***Rhodococcus* (subgen. *Anisorhodococcus*) *rhodnii* comb. nov.**<br>(rhod'ni.i. N.L. masc. n. *Rhodnius*, generic name of the reduvid bud; N.L. gen. masc. n. *rhodnii*, of *Rhodnius*) | **Basonym:** *Rhodococcus rhodnii* Goodfellow and Alderson 1979 (Approved Lists 1980) | The description of this taxon is as given by Goodfellow and Alderson [57] and Jones and Goodfellow [54]. The type strain has a G+C content of 69.5 % and circular genome (assembly) with a size of ≈4.4 Mbp.<br>**Type strain:** N445<sup>T</sup> (= ATCC 35071<sup>T</sup> = CIP 104181<sup>T</sup> = DSM 43336<sup>T</sup> = IEGM 555<sup>T</sup> = JCM 3203<sup>T</sup> = KCC A-203<sup>T</sup> = LMG 5363<sup>T</sup> = NBRC 100604<sup>T</sup> = NCIB 11279<sup>T</sup> = NCIMB 11279<sup>T</sup> = NRRL B-16535<sup>T</sup> = VKM Ac-1187<sup>T</sup>). |
| ***Rhodococcus* (subgen. *Anisorhodococcus*) *triatomae* comb. nov.**<br>(tri.a.to'mae. N.L. gen. fem. n. *triatomae*, of *Triatoma*, a zoological genus of blood-sucking bug from which the micro-organism was isolated) | **Basonym:** *Rhodococcus triatomae* Yassin 2005 | The description of this taxon is as given by Yassin [71]. The type strain has a G+C content of 69.5 % and circular genome with a size of ≈4.7 Mbp.<br>**Type strain:** IMMIB RIV-85<sup>T</sup> (= CCUG 50202<sup>T</sup> = DSM 44892<sup>T</sup> = NBRC 103116<sup>T</sup>). |
| ***Rhodococcus* (subgen. *Anisorhodococcus*) *wratislaviensis* comb. nov.**<br>(wra.tis.la.vi.en'sis. M.L. n. *wratislavia*, Wroclaw, Poland; N.L. masc./fem. adj. *wratislaviensis*, pertaining to Wroclaw) | **Basonyms:** *Rhodococcus wratislaviensis* (Goodfellow *et al.* 1995) Goodfellow *et al.* 2002<br>**Synonym:** *Tsukamurella wratislaviensis* Goodfellow *et al.* 1995 | The description of this taxon is as given by Goodfellow *et al.* [72] and Jones and Goodfellow 2012 [54]. The type strain has a G+C content of 67.5 % and genome (assembly) size of ≈7.7 Mbp.<br>**Type strain:** N805<sup>T</sup> (= ATCC 51786<sup>T</sup> = CCUG 38518<sup>T</sup> = CIP 105033<sup>T</sup> = DSM 44107<sup>T</sup> = JCM 9689<sup>T</sup> = NBRC 100605<sup>T</sup> = NCIMB 13082<sup>T</sup> = VKM Ac-1986<sup>T</sup>). |
| ***Rhodococcus* (subgen. *Pararhodococcus*) *maanshanensis* comb. nov.**<br>(maan.shan.en'sis. N.L. masc./fem. adj. *maanshanensis*, of or belonging to Maanshan, the source of the soil from which the organism was isolated) | **Basonym:** *Rhodococcus maanshanensis* Zhang *et al.* 2002 | The description of this taxon is as given by Zhang *et al.* [73]. The type strain has a G+C content of 69.2 % and a genome (assembly) size of $\approx 5.7$ Mbp.<br>**Type strain:** M712<sup>T</sup> (= CGMCC 4.1720<sup>T</sup> = DSM 44675<sup>T</sup> = JCM 11374<sup>T</sup> = NBRC 100610<sup>T</sup>). |
| ***Rhodococcus* (subgen. *Pararhodococcus*) *olei* comb. nov.**<br>(o'le.i. L. gen. neut. n. *olei*, of/from oil, as the organism was isolated from oil-contaminated soil) | **Basonym:** *Rhodococcus olei* Chaudhary and Kim 2018 | The description of this taxon is as given by Chaudhary and Kim [74]. The type strain has a G+C content of 70 % and a genome size of $\approx 6.2$ Mbp.<br>**Type strain:** Ktm-20<sup>T</sup> (=KEMB 9005-695<sup>T</sup>=KACC 19390<sup>T</sup>=JCM 32206<sup>T</sup>). |
| ***Rhodococcus* (subgen. *Pararhodococcus*) *oryzae* comb. nov.**<br>(o.ry'zae. L. gen. fem. n. *oryzae*, of rice) | **Basonym:** *Rhodococcus oryzae* Li *et al.* 2020 | The description of this taxon is as given by Li *et al.* [75]. The type strain has a G+C content of 69.2 % and a genome size of $\approx 5.4$ Mbp.<br>**Type strain:** NEAU CX67<sup>T</sup> (= CCTCC AB 2018233<sup>T</sup> = DSM 107701<sup>T</sup>). |
| ***Rhodococcus* (subgen. *Pararhodococcus*) *spelaei* comb. nov.**<br>(spe'lae.i. L. gen. neut. n. *spelaei*, of cave, referring to the site which the type strain was isolated) | **Basonym:** *Rhodococcus spelaei* Lee and Kim 2021 | The description of this taxon is as given by Lee and Kim [76]. The type strain has a G+C content of 69.2 % and a genome size of $\approx 4.8$ Mbp.<br>**Type strain:** C9-5<sup>T</sup> (= DSM 107558<sup>T</sup> = KACC 19822<sup>T</sup>). |
| ***Rhodococcus* (subgen. *Pararhodococcus*) *tukisamuensis* comb. nov.**<br>(tu.ki.sa.mu.en'sis. N.L. masc./fem. adj. *tukisamuensis*, referring to Tukisamu, a town in Sapporo, Hokkaido, Japan, where the type strain was isolated) | **Basonym:** *Rhodococcus tukisamuensis* Matsuyama *et al.* 2003 | The description of this taxon is as given by Matsuyama *et al.* [77]. The type strain has a G+C content of 69.8 % and a genome size of $\approx 5.5$ Mbp.<br>**Type strain:** Mb8<sup>T</sup> (= DSM 44783<sup>T</sup> = JCM 11308<sup>T</sup> = NBRC 100609<sup>T</sup> = NCIMB 13903<sup>T</sup>). |
| ***Rhodococcus* (subgen. *Rhodococcus* Zopf 1891) *aetherivorans* comb. nov.**<br>(ae.the.ri.vorans. N.L. *aetherum*, ether; L. pres. part. *vorans*, eating, devouring; N.L. masc. part. adj. *aetherivorans*, devouring ether) | **Basonym:** *Rhodococcus aetherivorans* Goodfellow *et al.* 2004 | The description of this taxon is as given by Zhang *et al.* [78]. The type strain has a G+C content of 70.5% and circular genome with a size of $\approx 6.4$ Mbp.<br>**Type strain:** 10bc312<sup>T</sup> (= DSM 44752<sup>T</sup> = JCM 14343<sup>T</sup> = NCIMB 13964<sup>T</sup>). |
| ***Rhodococcus* (subgen. *Rhodococcus* Zopf 1891) *artemisiae* comb. nov.**<br>(ar.te.mi'si.ae. L. fem. n. *artemisia*, mugwort, also a plant genus; L. gen. fem. n. *artemisiae*, of Artemisia, isolated from Artemisia annua L.) | **Basonym:** *Rhodococcus artemisiae* Zhao *et al.* 2012 | The description of this taxon is as given by Zhao *et al.* [79]. The type strain has a G+C content of 65.5% and circular genome with a size of $\approx 7.1$ Mbp.<br>**Type strain:** YIM 65754<sup>T</sup> (= CCTCC AA 209042<sup>T</sup> = DSM 45380<sup>T</sup>). |
| ***Rhodococcus* (subgen. *Rhodococcus* Zopf 1891) *chondri* comb. nov.**<br>(chon'dri. N.L. gen. masc. n. *chondri*, of the algal genus Chondrus) | **Basonym:** *Rhodococcus chondri* Girão *et al.* 2024 | The description of this taxon is as given by Girão *et al.* [80]. The type strain has a G+C content of 67% and circular genome with a size of $\approx 5.3$ Mbp.<br>**Type strain:** CC-R104<sup>T</sup> (= LMG 33233<sup>T</sup> = UCCCB 171<sup>T</sup>). |
| ***Rhodococcus* (subgen. *Rhodococcus* Zopf 1891) *coprophilus* comb. nov.**<br>(co.pro'phi.lus. Gr. fem. n. *kopros*, dung, faeces; N.L. adj. *philus* -a -um, friend, loving; from Gr. adj. *philos* -ê -on, loving; N.L. masc. adj. *coprophilus*, dung loving) | **Basonym:** *Rhodococcus coprophilus* Rowbotham and Cross 1979 (Approved Lists 1980) | The description of this taxon is as given by Rowbotham and Cross [81]. The type strain has a G+C content of 67% and circular genome with a size of $\approx 4.5$ Mbp.<br>**Type strain:** ATCC 29080<sup>T</sup> (= CIP 104178<sup>T</sup> = CUB 687<sup>T</sup> = DSM 43347<sup>T</sup> = IEGM 600<sup>T</sup> = JCM 3200<sup>T</sup> = LMG 5357<sup>T</sup> = NBRC 100603<sup>T</sup> = NCIB 11211<sup>T</sup> = NCIMB 11211<sup>T</sup> = NCTC 10994<sup>T</sup> = NRRL B-16537<sup>T</sup> = VKM Ac-571<sup>T</sup>). |
| ***Rhodococcus* (subgen. *Rhodococcus* Zopf 1891) *gordoniae* comb. nov.**<br>(gor.don'i.ae. N.L. gen. fem. n. *gordoniae*, of Gordon, named after Ruth Gordon, a celebrated microbial systematist) | **Basonym:** *Rhodococcus gordoniae* Jones *et al.* 2004 | The description of this taxon is as given in Jones *et al.* [82]. The type strain has a G+C content of 68% and circular genome with a size of $\approx 4.9$ Mbp.<br>**Type strain:** W 4937<sup>T</sup> (= DSM 44689<sup>T</sup> = JCM 12658<sup>T</sup> = NCTC 13296<sup>T</sup>). |
| ***Rhodococcus* (subgen. *Rhodococcus* Zopf 1891) *indonesiensis* comb. nov.**<br>(in.do.ne.si.en'sis. N.L. masc. adj. *indonesiensis*, of or pertaining to Indonesia, the source of the isolate) | **Basonym:** *Rhodococcus indonesiensis* Kusuma *et al.* 2024 | The description of this taxon is as given in Kusuma *et al.* [83]. The type strain has a G+C content of 70% and circular genome with a size of $\approx 5.5$ Mbp.<br>**Type strain:** CSLK01-03<sup>T</sup> (= CCMM B1310<sup>T</sup> = CSLK01-03<sup>T</sup> = ICEBB 6<sup>T</sup> = NCIMB 15214<sup>T</sup>). |
| ***Rhodococcus* (subgen. *Rhodococcus* Zopf 1891) *phenolicus* comb. nov.**<br>(phen.o'li.cus. N.L. neut. n. *phenol*, common name for industrial solvent hydroxybenzene; L. adj. suff. -icus -a -um, suffix used with the sense of belonging to; N.L. masc. adj. *phenolicus*, pertaining to phenol) | **Basonym:** *Rhodococcus phenolicus* Rehfuss and Urban 2006 | The description of this taxon is as given by Rehfuss and Urban. [84]. The type strain has a G+C content of 68.5% and circular genome with a size of $\approx 6.3$ Mbp.<br>**Type strain:** G2P<sup>T</sup> (= DSM 44812<sup>T</sup> = JCM 14914<sup>T</sup> = NRRL B-24323<sup>T</sup>). |
| ***Rhodococcus* (subgen. *Rhodococcus* Zopf 1891) *pyridinivorans* comb. nov.**<br>(py.ri.di.ni.vo.rans. N.L. neut. n. *pyridinum*, pyridine; L. inf. v. *vorare*, to devour; N.L. masc. part. adj. *pyridinivorans*, pyridine-devouring) | **Basonym:** *Rhodococcus pyridinivorans* Yoon *et al.* 2000<br>**Synonym:** *Rhodococcus biphenylivorans* Su *et al.* 2015 | The description of this taxon is as given in Yoon *et al.* [85]. The type strain has a G+C content of 68% and circular genome with a size of $\approx 5.3$ Mbp.<br>**Type strain:** PDB9<sup>T</sup> (= DSM 44555<sup>T</sup> = JCM 10940<sup>T</sup> = KCCM 80005<sup>T</sup> = KCTC 0647BP<sup>T</sup> = NBRC 100608<sup>T</sup>). |
| ***Rhodococcus* (subgen. *Rhodococcus* Zopf 1891) *rhodochrous* comb. nov.**<br>(rho.do.chro.us. N.L. masc. adj. *rhodochrous*, rose colored) | **Basonym:** "Staphylococcus rhodochrous" Zopf 1891<br>**Synonyms:** *Rhodococcus roseus* (ex Grotenfelt 1889) Tsukamura *et al.* 1991; *Rhodococcus rhodochrous* (Zopf 1891) Tsukamura 1974 (Approved Lists 1980) | The description of this taxon is as given by Tsukamura [86], Goodfellow and Alderson [57], and Jones and Goodfellow [54]. The type strain has a G+C content of 68% and circular genome with a size of $\approx 5.2$ Mbp.<br>**Type strain:** ATCC 13808<sup>T</sup> (= CCUG 47165<sup>T</sup> = CIP 104376<sup>T</sup> = HAMBI 1959<sup>T</sup> = IEGM 62<sup>T</sup> = IFO 16069<sup>T</sup> = JCM 3202<sup>T</sup> = LMG 5365<sup>T</sup> = NBRC 16069<sup>T</sup> = NCTC 10210<sup>T</sup> = NRRL B-16536<sup>T</sup> = NRRL B-2149<sup>T</sup> = VKM Ac-1227<sup>T</sup>). |
| ***Rhodococcus* (subgen. *Rhodococcus* Zopf 1891) *ruber* comb. nov.**<br>(ru'ber. L. masc. adj. *ruber*, red) | **Basonym:** "*Streptothrix rubra*" Kruse 1896<br>**Synonyms:** *Rhodococcus ruber* (Kruse 1896) Goodfellow and Alderson 1977 (Approved Lists | The description of this taxon is as given by Goodfellow and Alderson [57], and Jones and Goodfellow [54]. The type strain |
| | 1980); *Rhodococcus electrodiphilus* Ramaprasad et al. 2018 | has a G+C content of 70.5% and circular genome with a size of $\approx 5.3$ Mbp.<br>**Type strain:** CIP 104180<sup>T</sup> (= DSM 43338<sup>T</sup> = IEGM 70<sup>T</sup> = IFO 15591<sup>T</sup> = JCM 3205<sup>T</sup> = KCC A-205<sup>T</sup> = LMG 5366<sup>T</sup> = NBRC 15591<sup>T</sup> = VKM Ac-1021<sup>T</sup>). |
| ***Rhodococcus* (subgen. *Rhodococcus* Zopf 1891) *yananensis* comb. nov.**<br>(yan.an.en'sis. N.L. masc. adj. *yananensis*, a city in Shaanxi Province, PR China, from where the type strain R was isolated) | **Basonym:** *Rhodococcus yananensis* Jiang et al. 2022 | The description of this taxon is as given by Jiang et al. [87]. The type strain has a G+C content of 68.5% and circular genome with a size of $\approx 4.2$ Mbp.<br>**Type strain:** FBM22-1<sup>T</sup> (= CCTCC AB 2020275<sup>T</sup> = FBM22-1<sup>T</sup> = KCTC 49502<sup>T</sup>). |
| ***Rhodococcus* (subgen. *Rhodococcus* Zopf 1891) *zopfii* comb. nov.**<br>(zopf'i.i. N.L. gen. masc. n. *zopfii*, of Zopf, named in honor of Wilhelm Friedrich Zopf, who described the bacterium *Rhodococcus rhodochrous*) | **Basonym:** *Rhodococcus zopfii* Stoecker et al. 1994 | The description of this taxon is as given by Stoecker et al. [88]. The type strain has a G+C content of 68% and circular genome with a size of $\approx 6.3$ Mbp.<br>**Type strain:** T1<sup>T</sup> (= ATCC 51349<sup>T</sup> = CIP 104275<sup>T</sup> = DSM 44108<sup>T</sup> = JCM 9919<sup>T</sup> = NBRC 100606<sup>T</sup> = NRRL B-16942<sup>T</sup>). |

The type species is *Rhodococcus equi* (Magnusson 1923) Goodfellow and Alderson 1977 (Approved Lists 1980).

### Pararhodococcus subgen. nov

(Pa.ra.rho.do.coc.cus. Gr. prep. *para*, beside; genuine; N.L. masc. n. *Rhodococcus* a genus name; N.L. masc. n. *Pararhodococcus*, beside *Rhodococcus,* most basal subgenus of the genus *Rhodococcus*).

Subgenus of genus *Rhodococcus* Zopf 1891 (Approved Lists 1980) emend. Val-Calvo and Vazquez-Boland 2023. Members share the same general morphological, physiological and chemotaxonomic characteristics of the genus *Rhodococcus* as described in Goodfellow and Alderson [57] and Jones and Goofellow [54]. Contains the species *Rhodococcus* (subgen. *Pararhodococcus*) *maanshanenensis*, *olei*, *oryzae*, *spelaei*, and *tukisamuensis,* isolated from soil in different ecosystems (mountainous, agricultural, cavern and urban). See **Table 1** for the description of the new name combinations. *Pararhodococcus* species have a genomic G+C content of 69 to 70% and size between 4.8 and 6.2 Mbp. They form the most basal monophyletic branch of the genus *Rhodococcus* Zopf 1891 (Approved Lists 1980) emend. Val-Calvo and Vazquez-Boland 2023 as determined by core-genome phylogenetic analysis. They can be differentiated from other rhodococci by means of relative evolutionary divergence (RED)-normalized tree clustering and genome relatedness index (GRI)-based network analysis, where they form an independent cluster or subnetwortk when applying clustering or graph cutoffs that isolate the *Prescottella* sublineage/subgenus as an independent cluster/subnetwork. In pairwise comparisons, the inter-subgenus AAI score of the current *Rhodococcus* (subgen. *Pararhodococcus*) circumscription is ≤73%.

Type species: *Rhodococcus* (subgen. *Pararhodococcus*) *maanshanensis*.

### *Anisorhodococcus* subgen. nov

(An.i.so.rho.do.coc.cus. Gr. masc. adj., unequal, dissimilar; N.L. masc. n. *Rhodococcus* a genus name; N.L. masc. n. *Anisorhodococcus,* unequal, dissimilar or uneven *Rhodococcus,* a subgenus of the genus *Rhodococcus*).

Subgenus of genus *Rhodococcus* Zopf 1891 (Approved Lists 1980) emend. Val-Calvo and Vazquez-Boland 2023. Members have the same general morphological, physiological and chemotaxonomic characteristics of the genus *Rhodococcus* as described in Goodfellow and Alderson [57] and Jones and Goofellow [54]. Subgenus *Anisorhodococcus* species contain the species *Rhodococcus* (subgen. *Anisorhodococcus*) *erythropolis*, *globerulus*, *jostii*, *koreensis*, *peudokoreensis*, *opacus*, *wratislaviensis, marinonascens, oxybenzonivorans, rhodnii*, and *triatomae*. See **Table 1** for the description of the new name combinations. They originate from diverse environments including soil, rhizosphere ecosystem, marine sediments and xenobiotic-contaminated sites, or are found as part of the gut microbiota of *Reduviidae* family triatomine hematophagous hemiptera (kissing bugs). The species of this heterogeneous subgenus are characterized by genomes of variable size and topology. Genome sizes range from ≈10 Mbp in the phenolic compound degraders *Rhodococcus* (*Anisorhodococcus*) *jostii* and *koreensis*, presumably caused by catabolic gene network amplification, to 4.5 and 4.7 Mbp for the insect gut symbionts *Rhodococcus* (subgen. *Anisorhodococcus*) *rhodnii* and *triatomae,* presumably due to host-adaptive reductive evolution. The species with larger genomes have linear chromosomes and several plasmids, from large (0.4 to 1 Mbp) invertron-like linear replicons to smaller circular plasmids; those with intermediate (e.g. *marinonascens*, 4.9 Mbp) and smaller (*rhodnii* and *triatomae*) genomes have circular chromosomes. G+C contents range between 64.5 and 69.5%. Members of subgenus *Rhodococcus* (subgen. *Anisorhodococcus*) correspond to *Rhodococcus* sublineage no. 4 described in this study. They can be distinguished from other *Rhodococcus* spp. by relative evolutionary divergence (RED)-normalized tree clustering and genome relatedness index (GRI)-based network analysis, where they form an independent cluster or subnetwortk when applying clustering or graph cutoffs that isolate the *Prescottella* sublineage/subgenus as an independent cluster/subnetwork (with the proviso that GRIs incorporating a measure of genome size in the divisor [POCP, AF] should be avoided because with these the genome size biases the clustering) [11]. In pairwise comparisons, the inter-subgenus AAI scores of the current *Rhodococcus* (subgen. *Anisorhodococcus*) circumscription are ≤73%.

The type species of the subgenus is *Rhodococcus* (subgen. *Anisorhodococcus*) *erythropolis*.

### *Rhodococcus* (Zopf 1891) subgen. nov

(Rho.do.coc.cus. Gr. neut. n. *rhodon*, the rose; N.L. masc. n. *coccus*, coccus; from Gr. masc. n. *kokkos*, grain, seed; N.L. masc. n. *Rhodococcus*, a red coccus) Subgenus of genus *Rhodococcus* Zopf 1891 (Approved Lists 1980) emend. Val-Calvo and Vazquez-Boland 2023 that contains the type species of the genus, *Rhodococcus rhodochrous*. Automatically formed in application of ICNP Rule 39 upon creation of the *Rhodococcus* subgenera *Prescottella*, *Anisorhodococcus*, and *Pararhodococcus*. The *Rhodococcus* (subgen. *Rhodococcus*) circumscription is comprised of the species *R. aeterivorans*, *R. artemisiae*, *R. chaundri*, *R. coprophilus*, *R. gordoniae*, *R. indonesiensis*, *R. phenolicus*, *R. pyridinivorans*, *R. rhodochrous*, *R. ruber*, *R. yananensis*, and *R. zopfii*. See **Table 1** for the description of the new name combinations.

### *Mycolicibacter* (Gupta *et al*. 2018) subgen. nov

(My.co.li.ci.bac’ter. N.L. neut. n. *acidum mycolicum*, mycolic acid; N.L. masc. n. *bacter*, rod; N.L. masc. n. *Mycolicibacter*, a genus of mycolic acid containing rod-shaped bacteria) The description of this taxon is as given by Gupta *et al.* 2018 [6]. Previously described as genus, it is lowered to subgenus of genus *Mycobacterium*. The type species is *Mycobacterium terrae* Wayne 1966 (Approved Lists 1980). The description of the new name combinations for the type species and the rest of the subgenus circumscription are provided in **Table 2**.

**Table 2.**
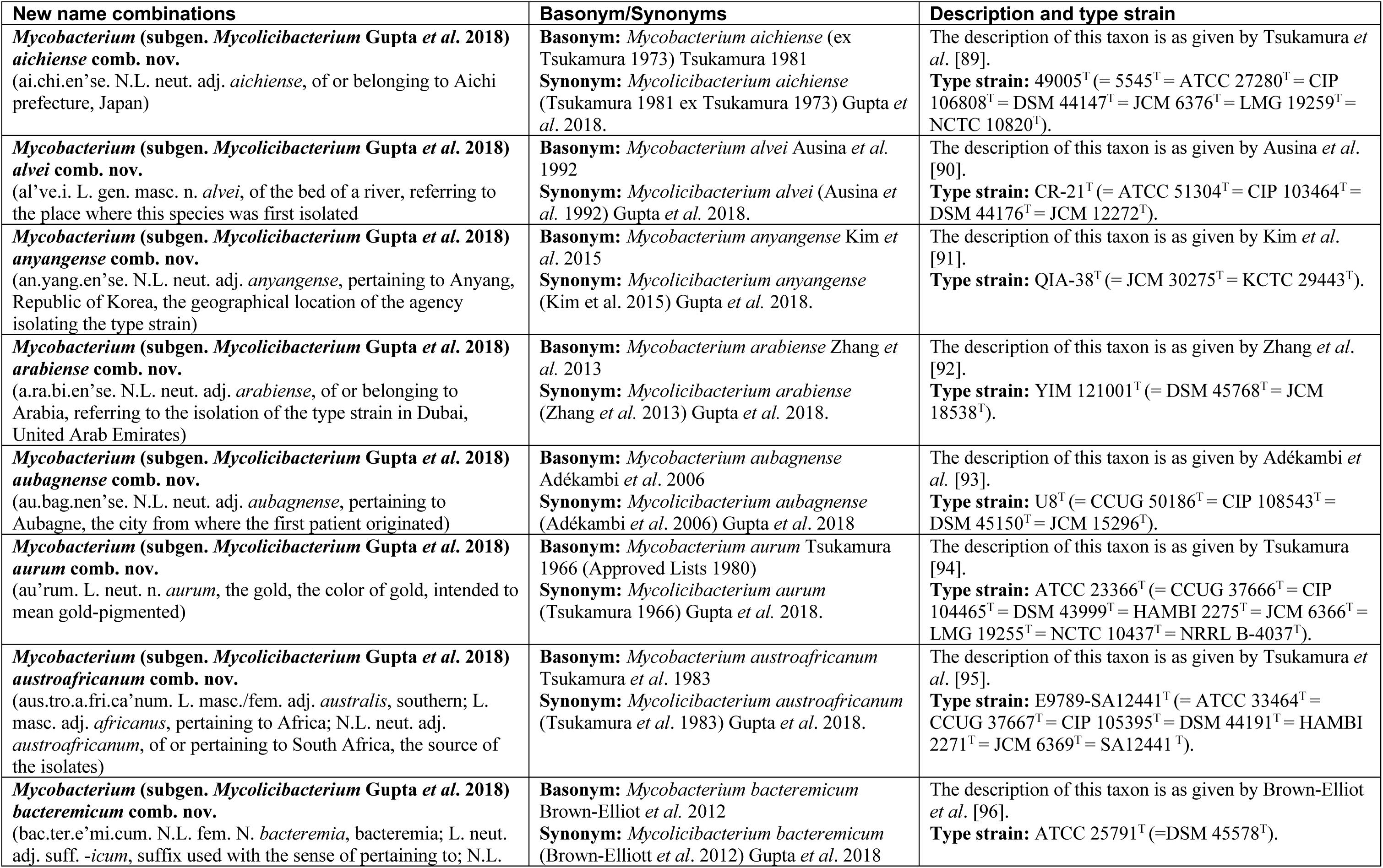
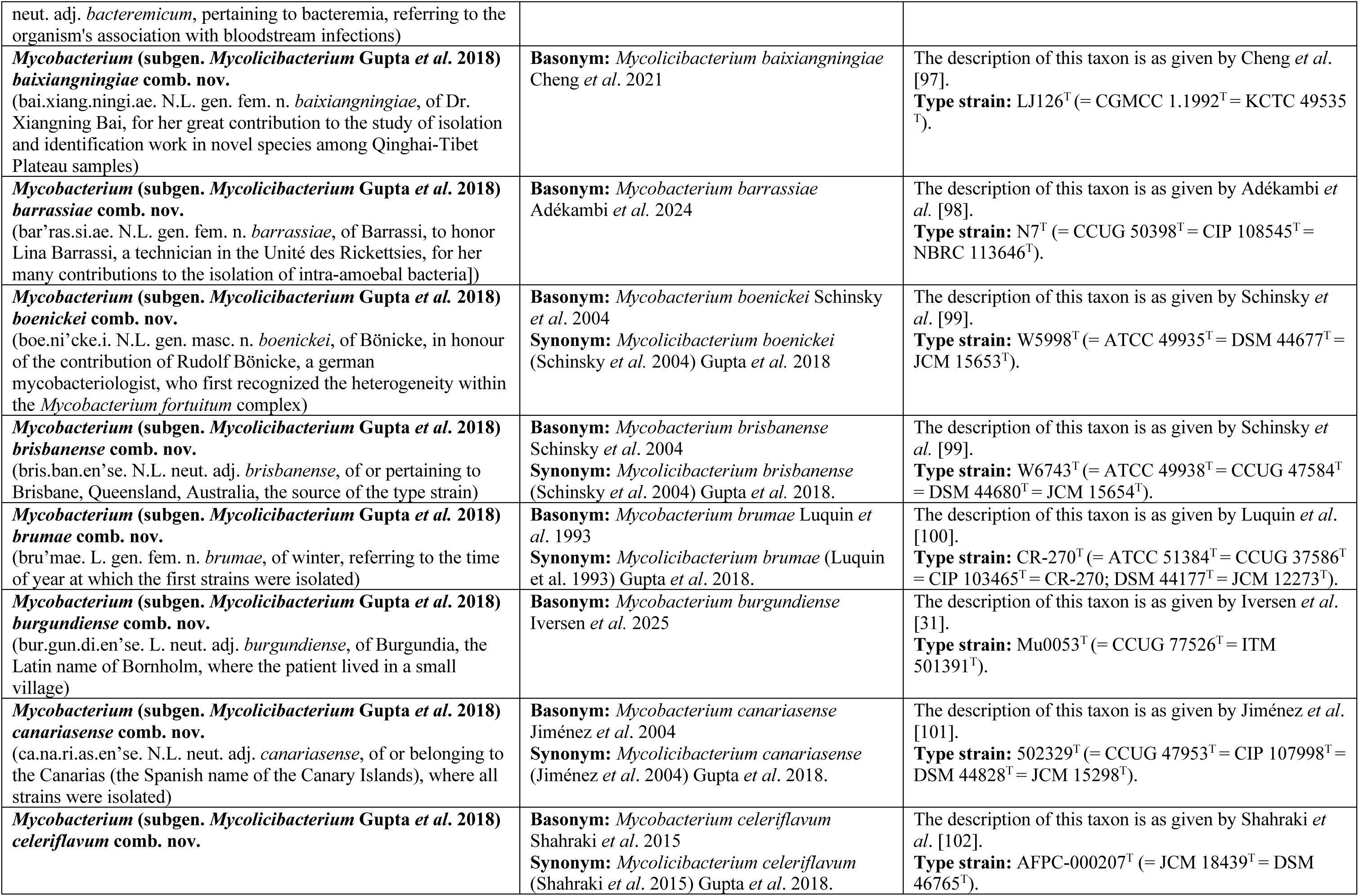
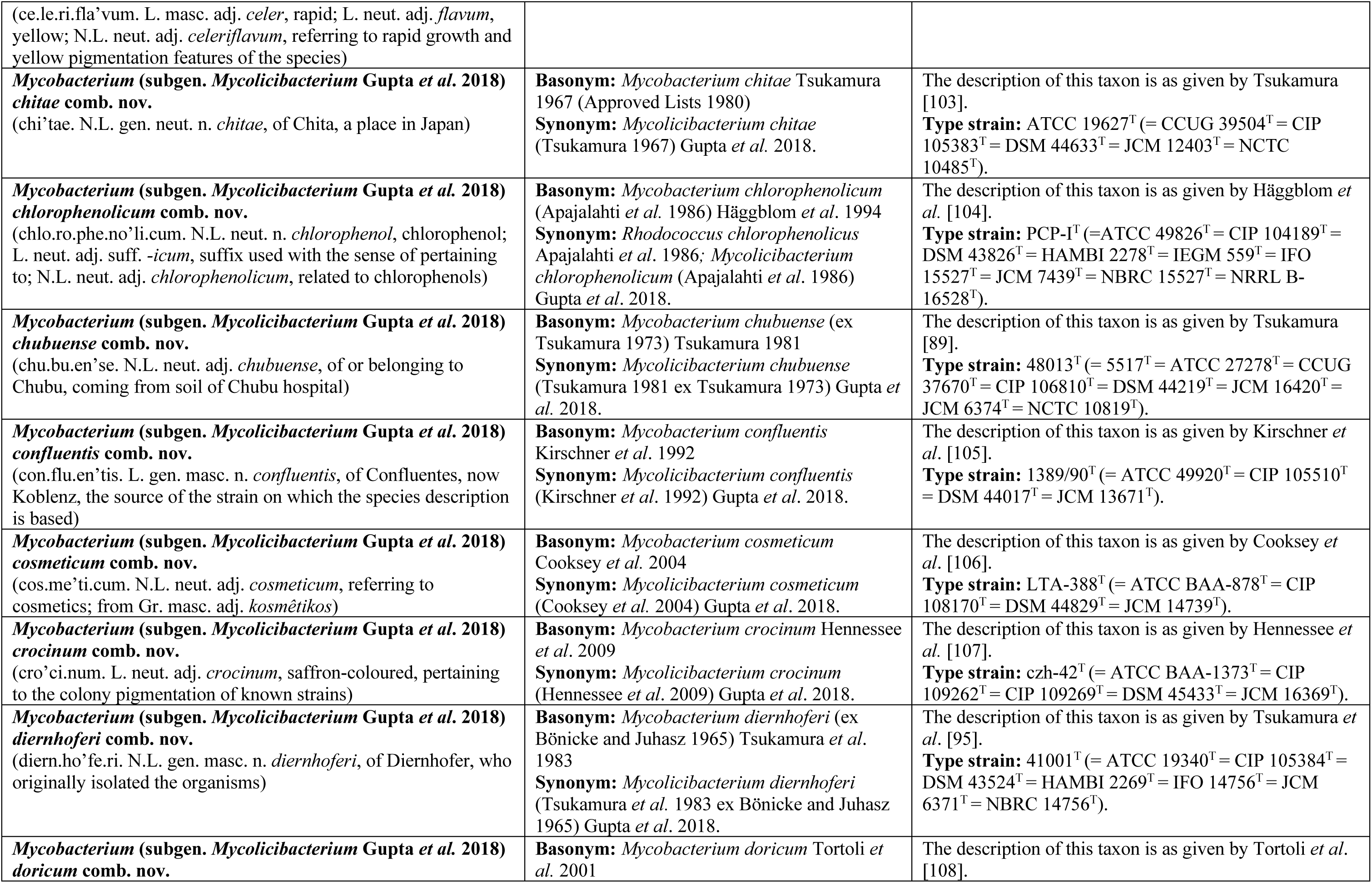
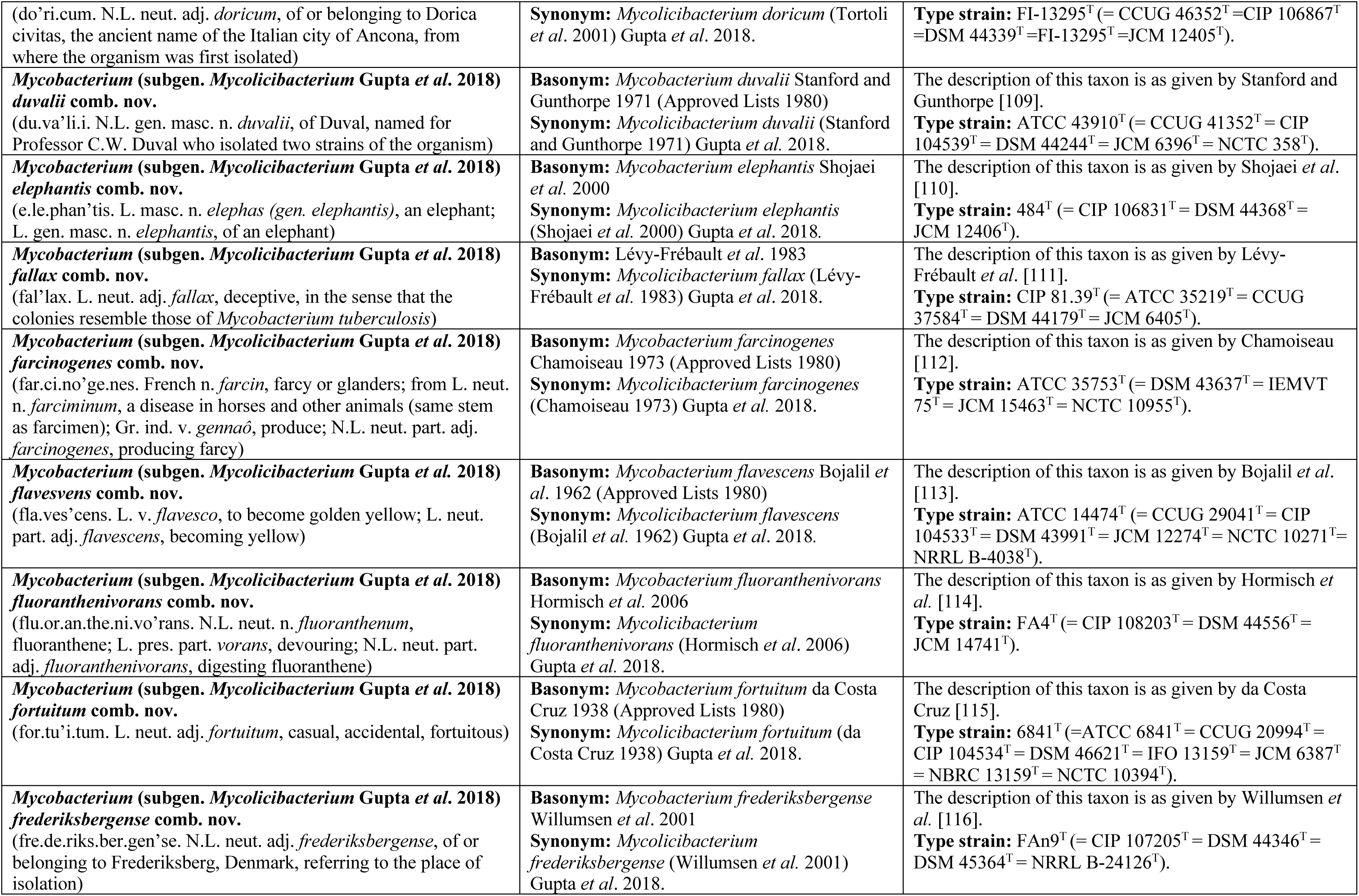
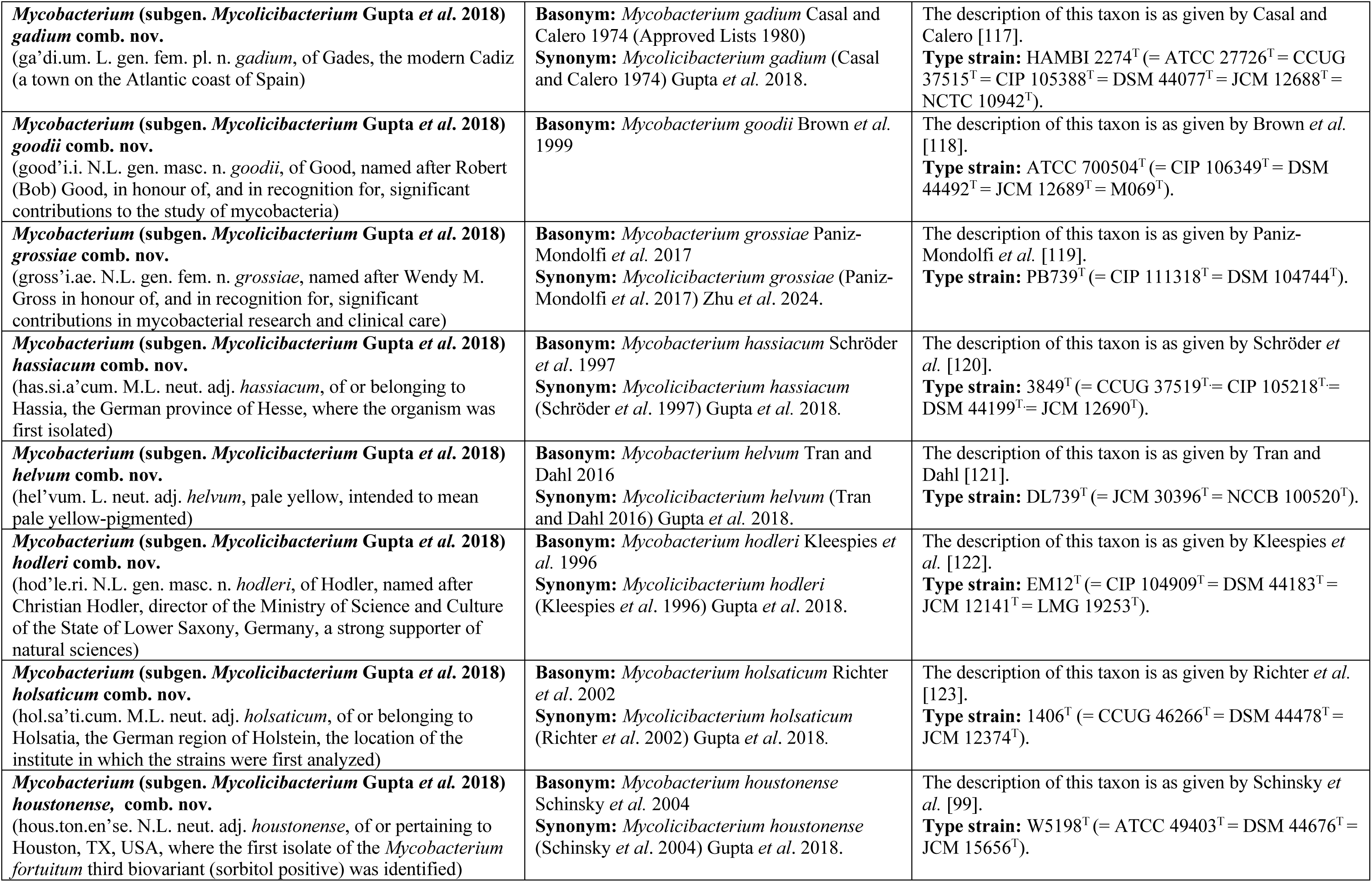
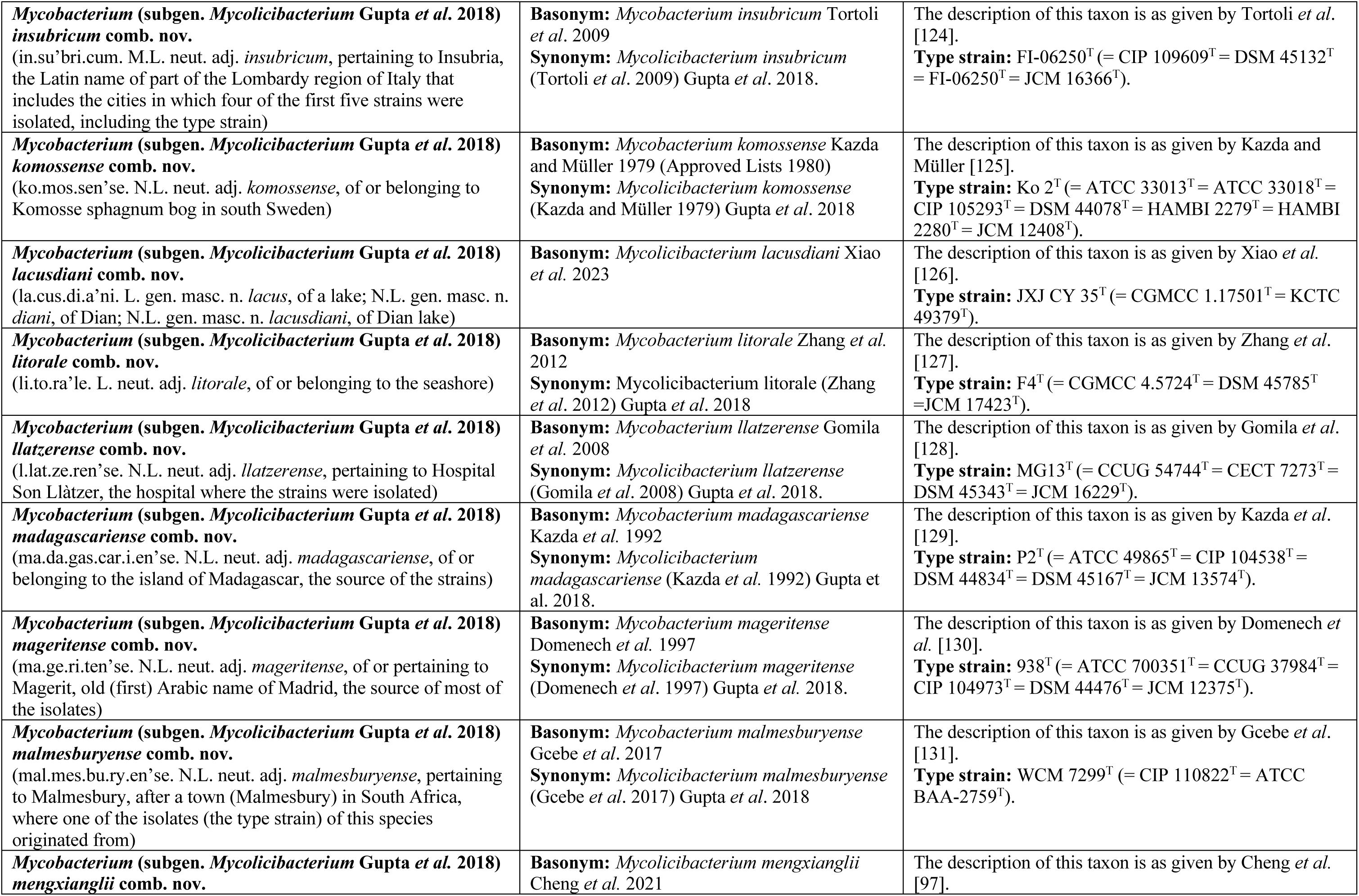
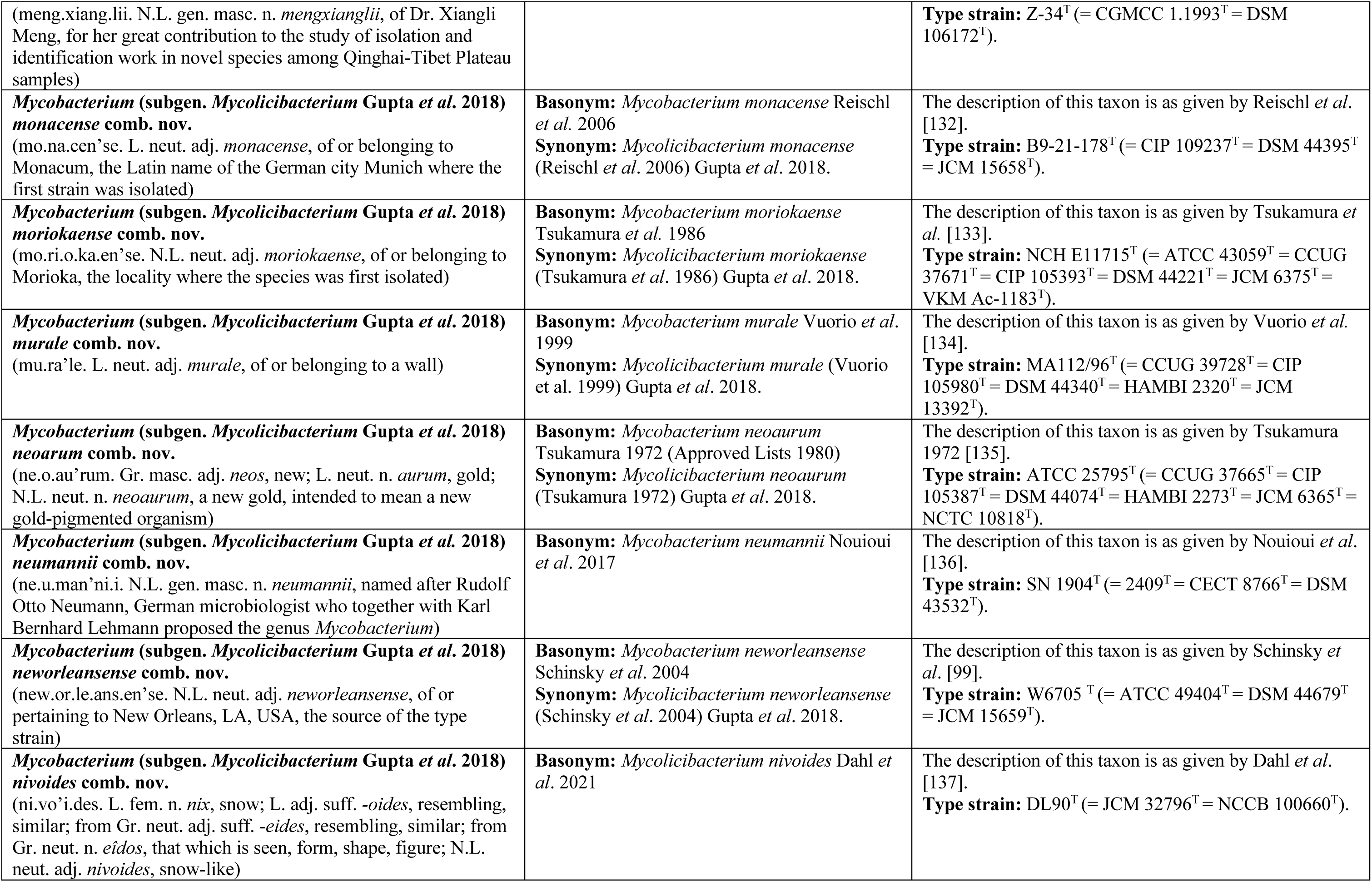
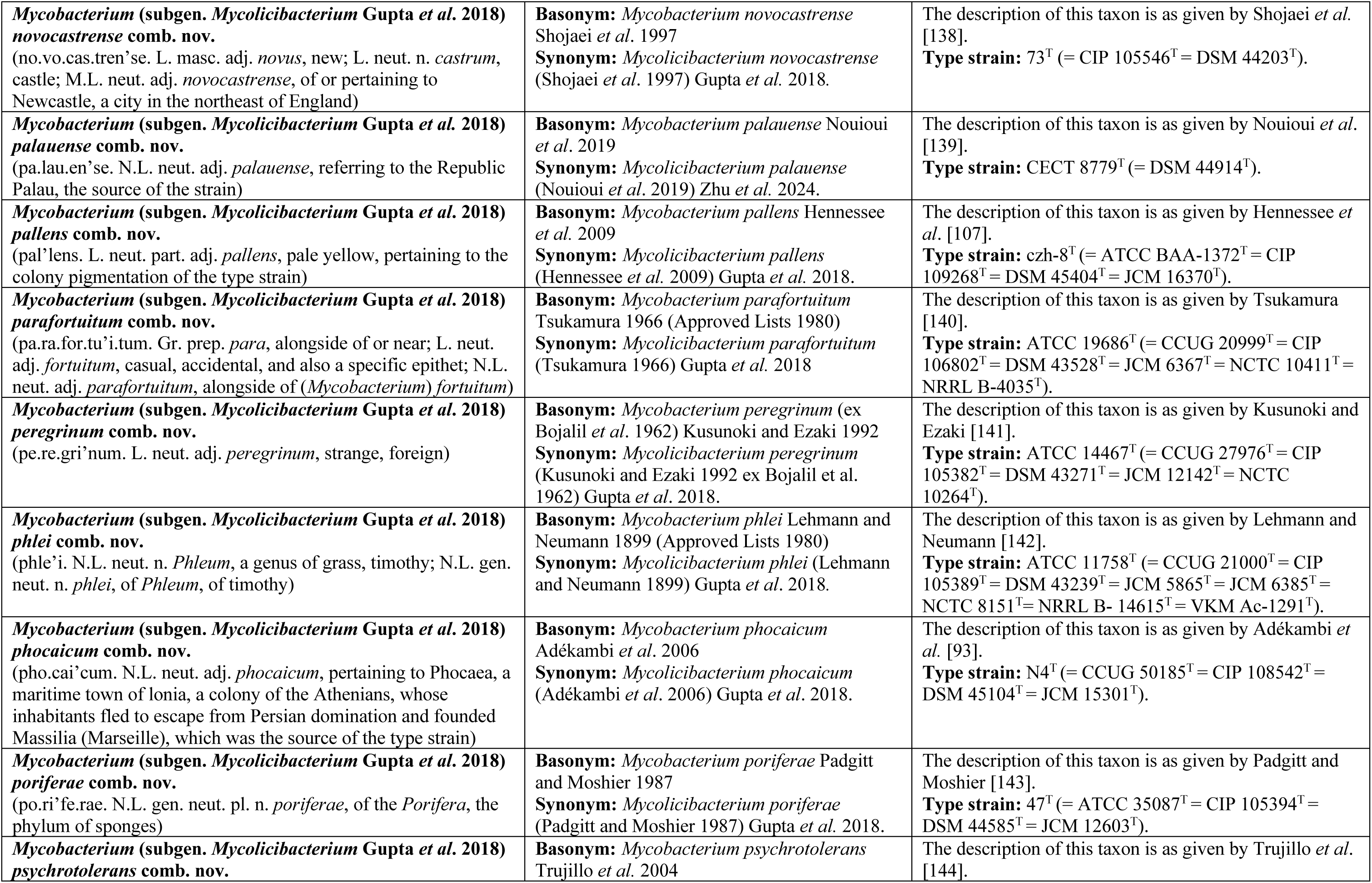
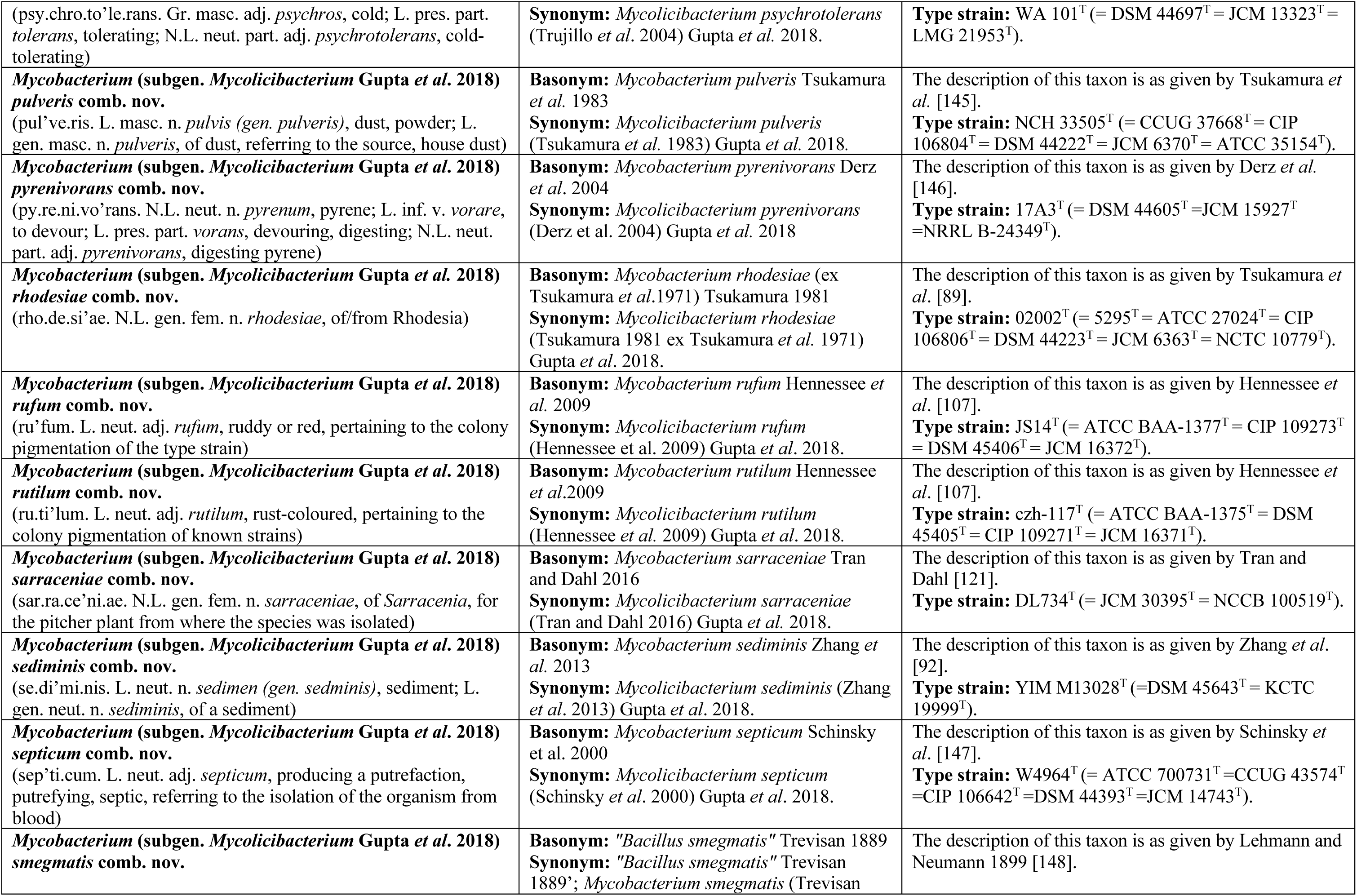
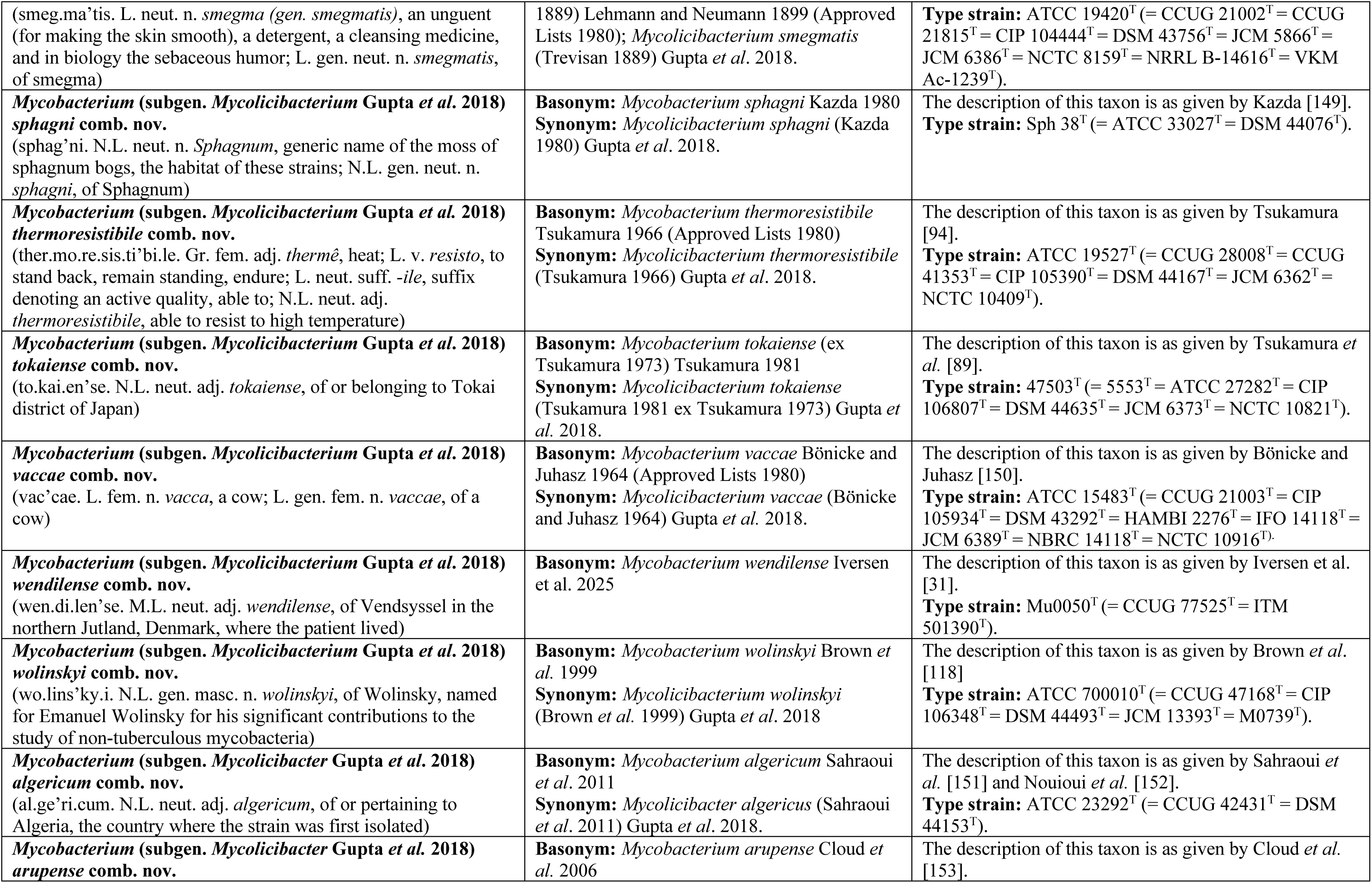
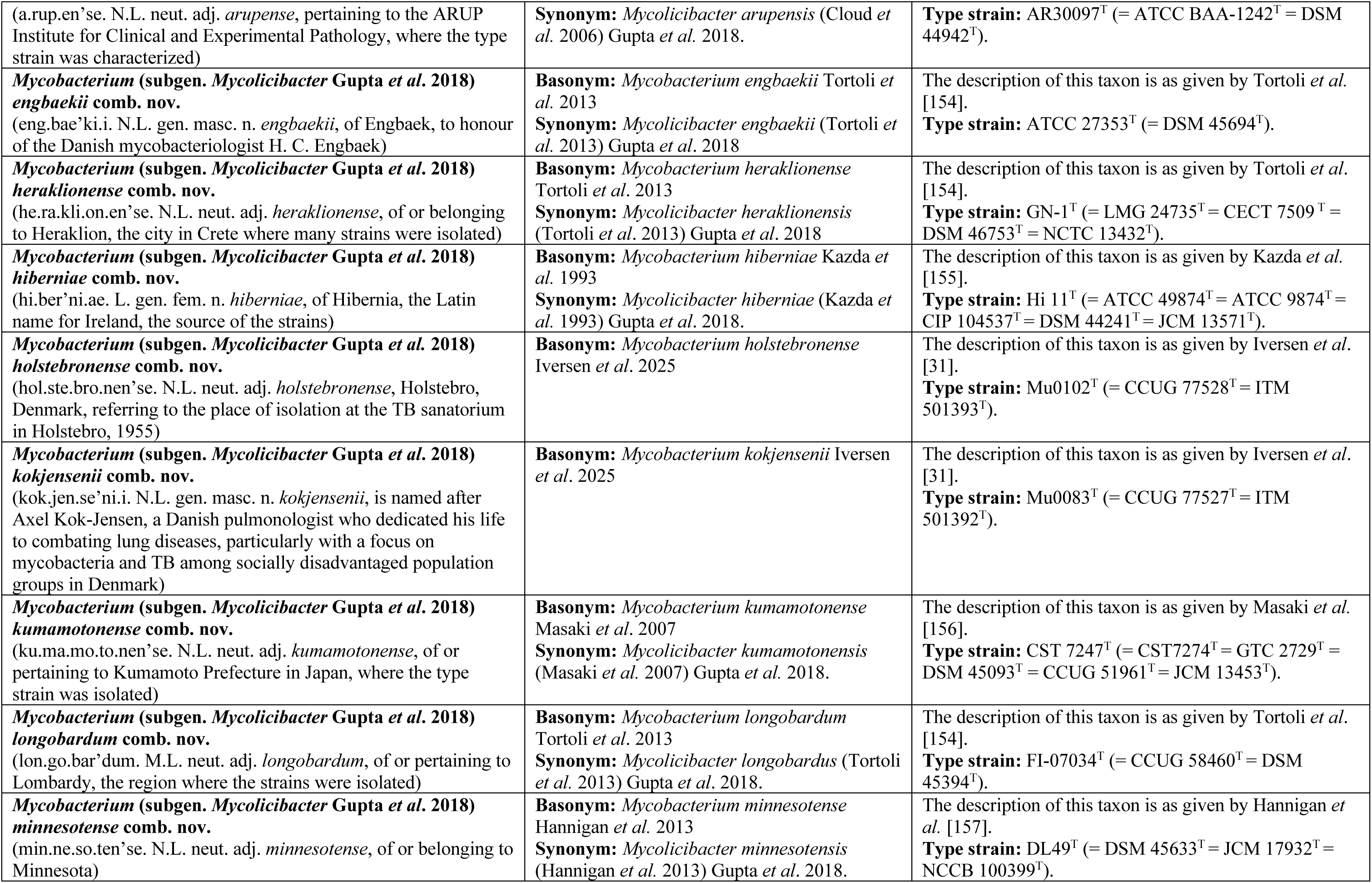
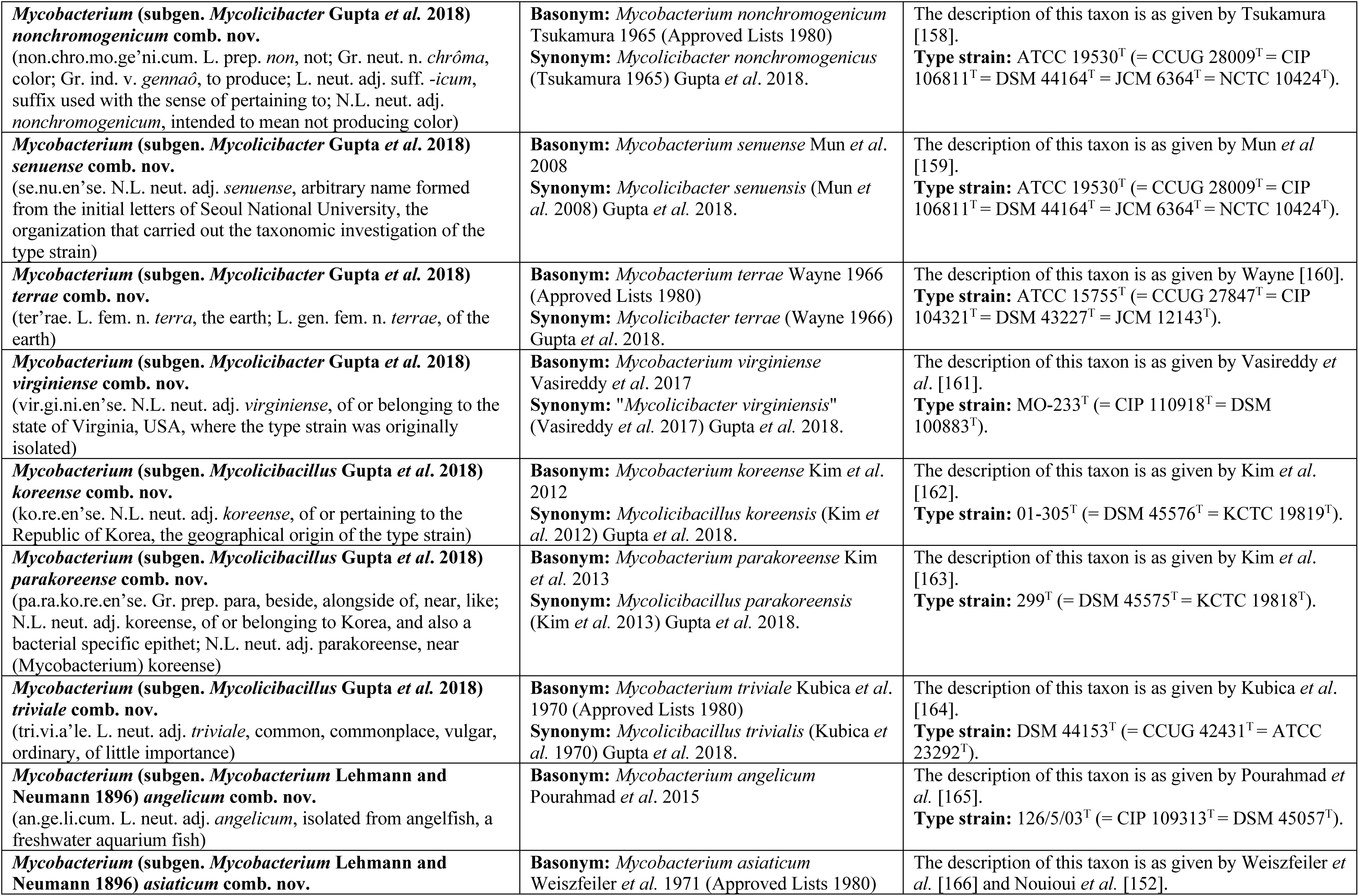
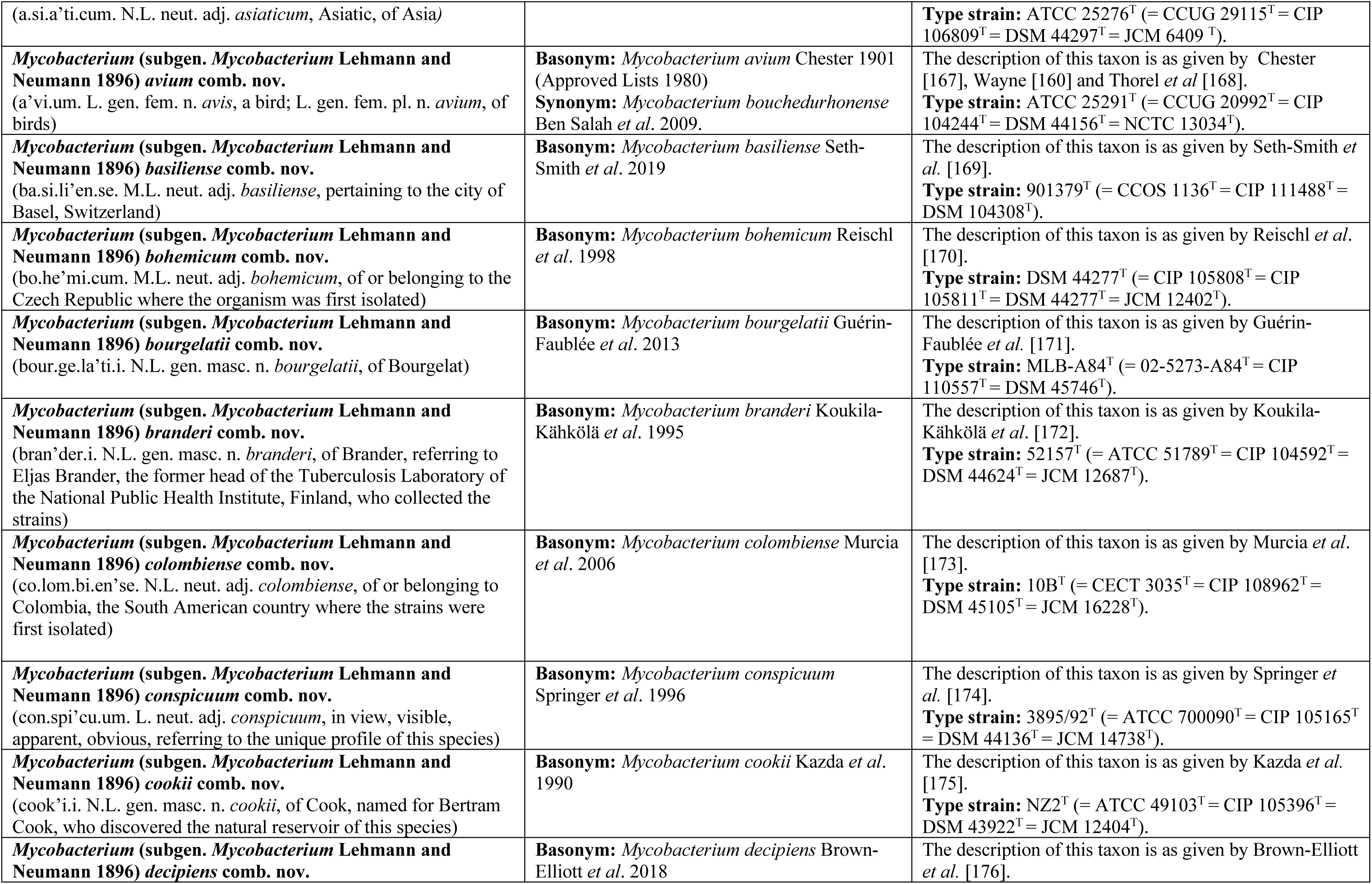
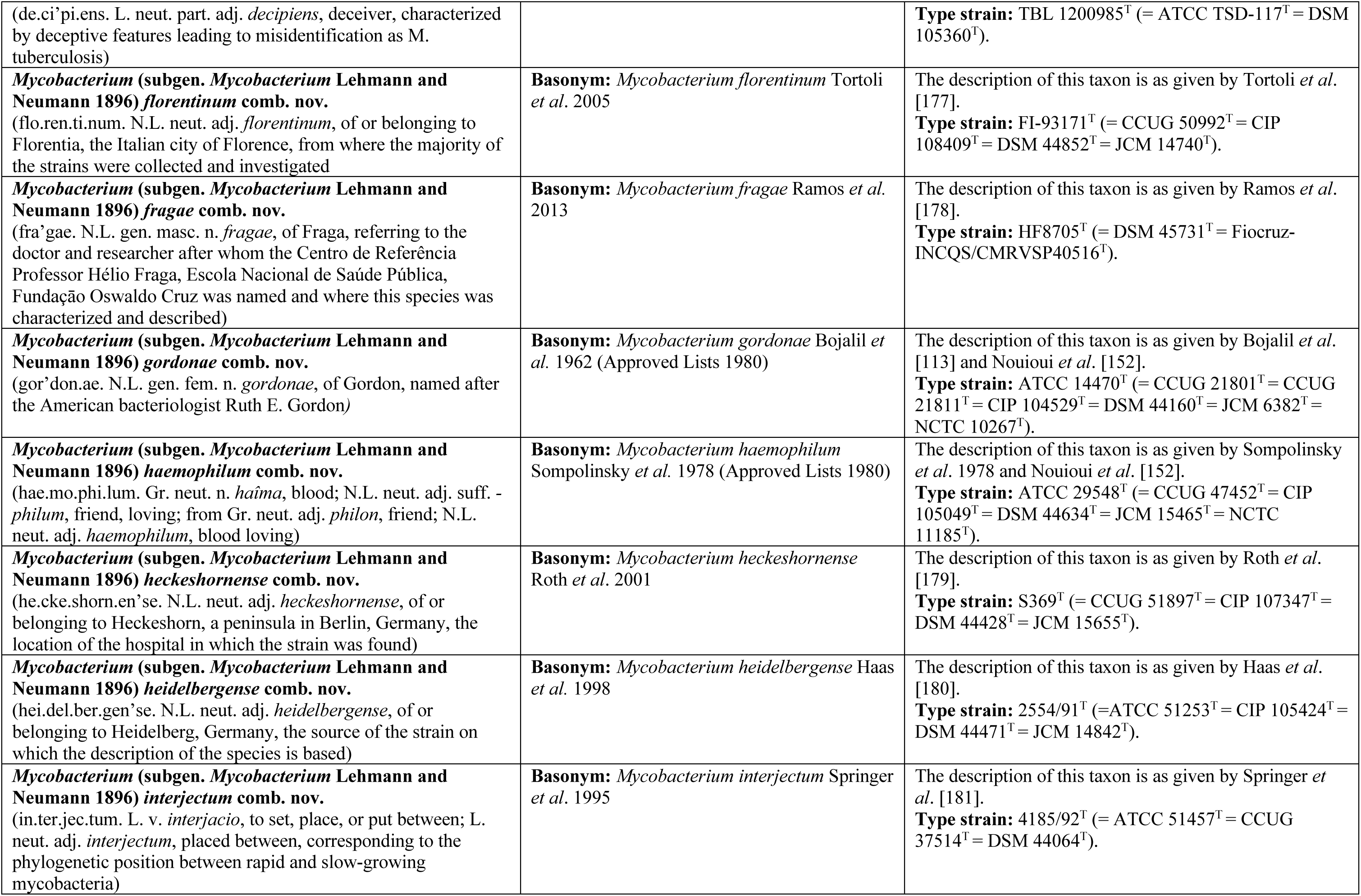
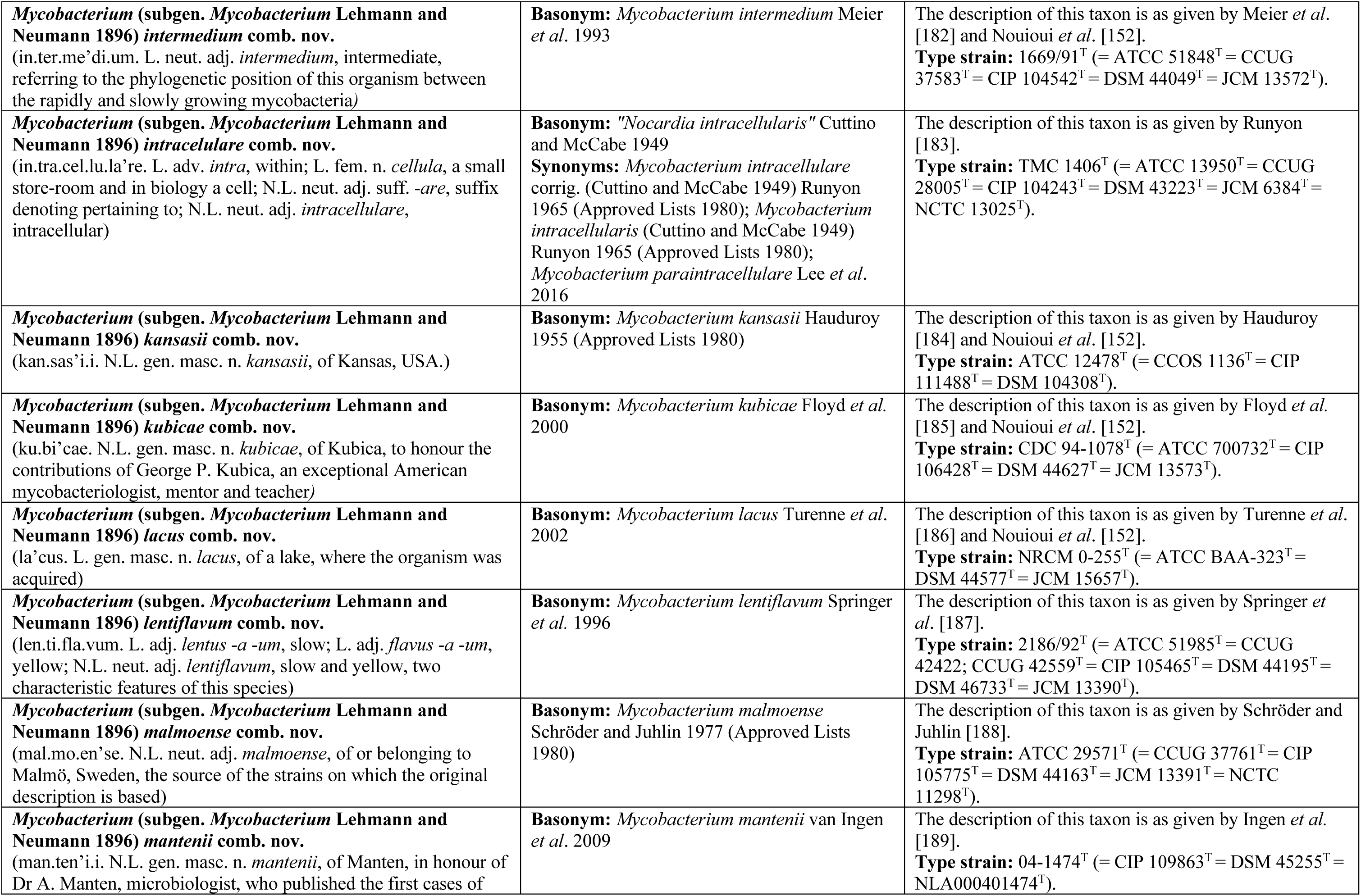
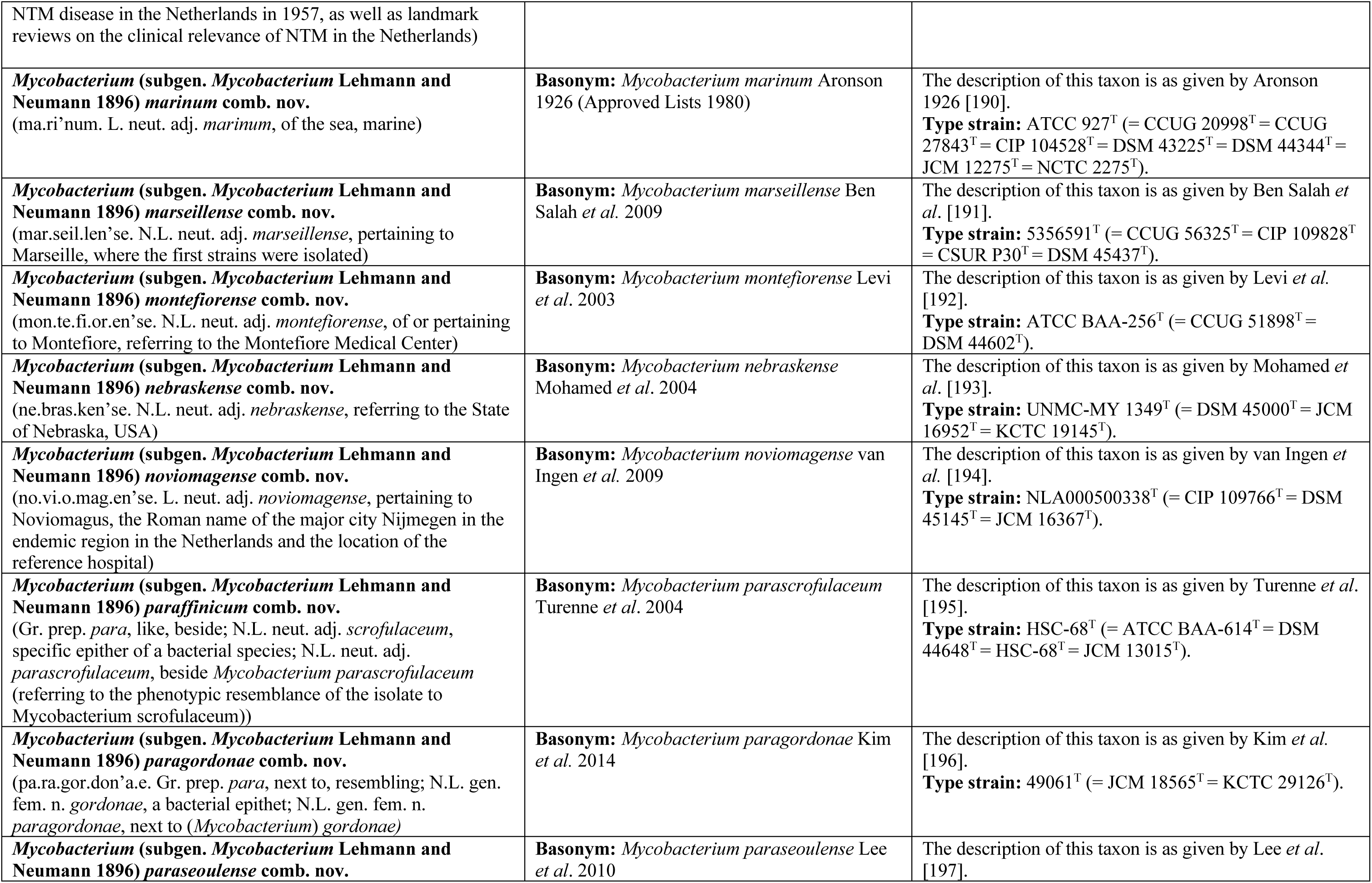
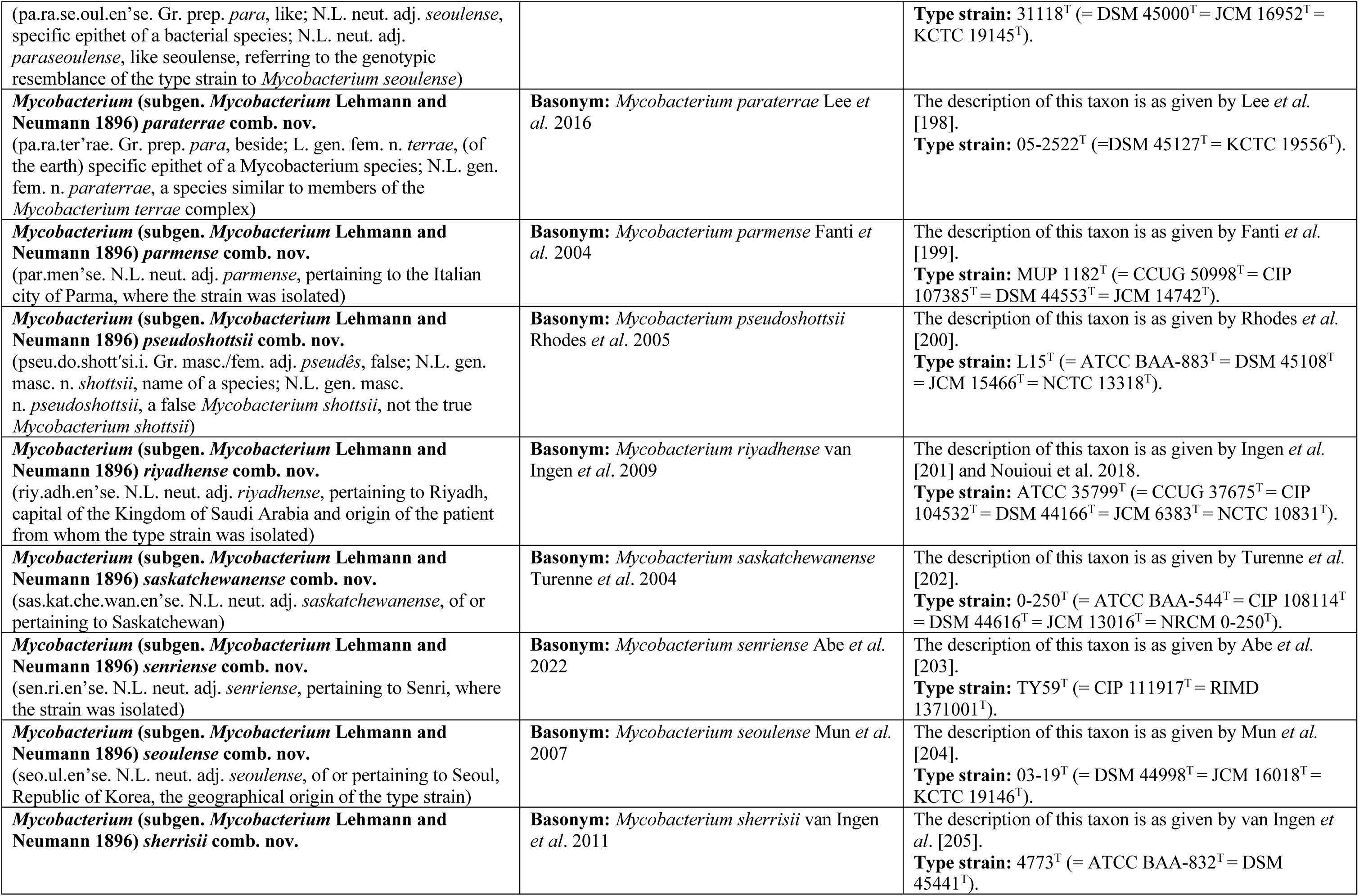
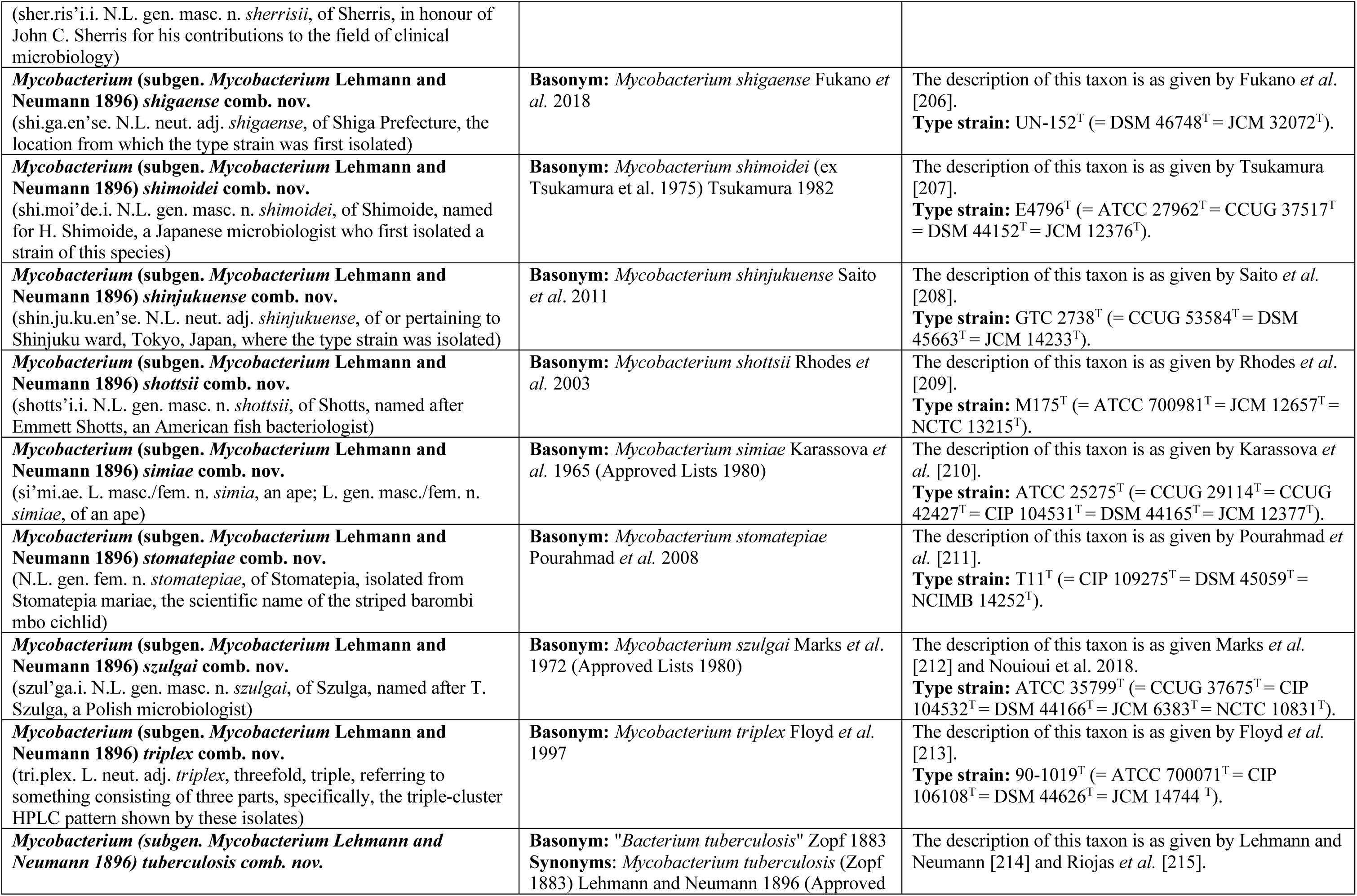
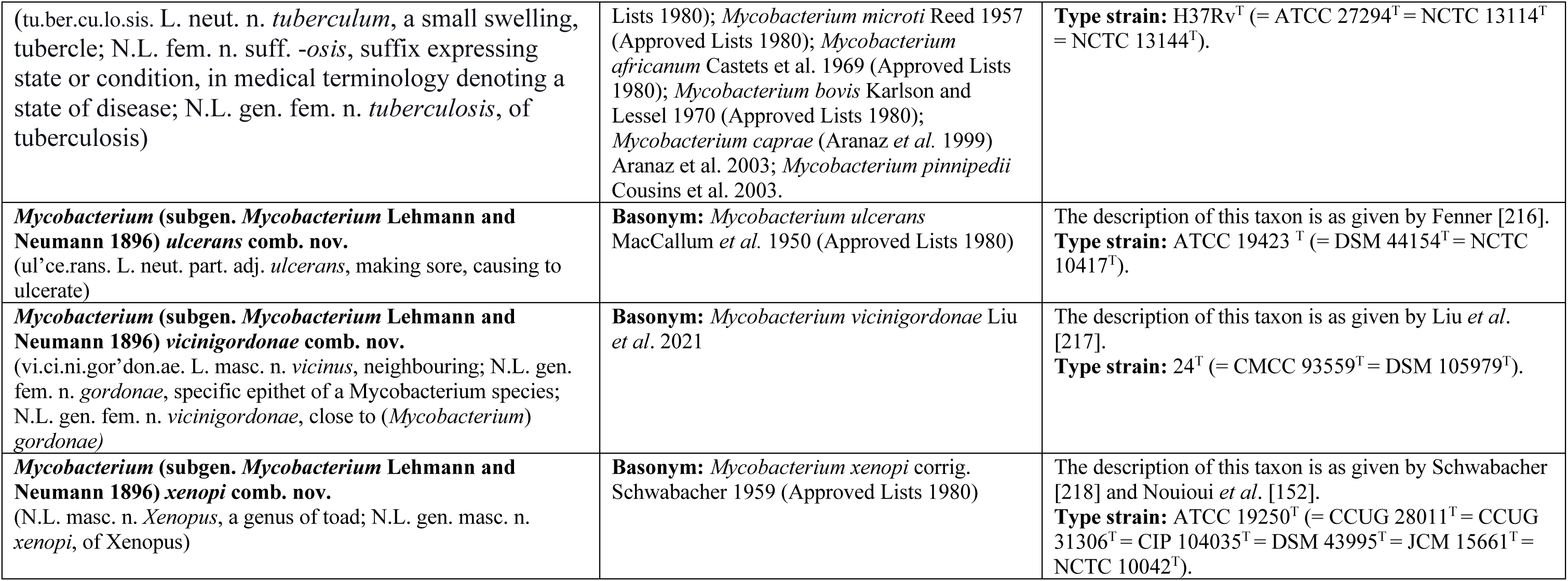
Descriptions of new *Rhodococcus* combinations.

### *Mycolicibacterium* (Gupta *et al*. 2018) subgen. nov

(My.co.li.ci.bac.te’ri.um. N.L. neut. n. *acidum mycolicum*, mycolic acid; N.L. neut. n. *bacterium*, a small rod; N.L. neut. n. *Mycolicibacterium*, a genus of mycolic acid containing rod-shaped bacteria). The description of this taxon is as given by Gupta *et al.* 2018 [6]. Previously described as genus, it is lowered to subgenus of genus *Mycobacterium*. The type species is *Mycobacterium fortuitum* da Costa Cruz 1938 (Approved Lists 1980). The description of the new name combinations for the type species and the rest of the subgenus circumscription are provided in **Table 2**.

### *Mycolicibacillus* (Gupta *et al*. 2018) subgen. nov

(My.co.li.ci.ba.cil’lus. N.L. neut. n. *acidum mycolicum*, mycolic acid; L. masc. n. *bacillus*, a small staff or rod; N.L. masc. n. *Mycolicibacillus*, a genus of mycolic acid containing rod-shaped bacteria).

The description of this taxon is as given by Gupta *et al.* [6]. Previously described as a genus, it is lowered to subgenus of genus *Mycobacterium*. The type species is *Mycobacterium triviale* Kubica *et al.* 1970 (Approved Lists 1980). The description of the new name combinations for the type species and the rest of the circumscription of the subgenus are provided in **Table 2**.

### *Mycobacterium* (Lehmann and Neumann 1896) subgen. nov

(My.co.bac.te’ri.um. Gr. masc. n. *mykês (gen. mykêtos)*, a mushroom, fungus; N.L. neut. n. *bacterium*, a rod; N.L. neut. n. *Mycobacterium*, a fungus rodlet) Subgenus of genus *Mycobacterium* Lehmann and Neumann 1896 (Approved Lists 1980) emend. Val-Calvo and Vázquez-Boland 2023 that contains the type species of the genus, *Mycobacterium tuberculosis* (Zopf 1883) Lehmann and Neumann 1896 (Approved Lists 1980). Automatically formed in application of ICNP Rule 39 upon creation of the *Mycobacterium* subgenera *Mycolicibacillus, Mycolicibacter,*and *Mycolicibacterium*. The description of the new name combinations for the type species and the rest of the circumscription of the subgenus are provided in **Table 2**.

### Rhodococcoides *navarretei* comb. nov

(na.var.re’te.i. N.L. gen. masc. n. navarretei, named in honour of Group Commander (AD) Eduardo Navarrete Pizarro of the Chilean Air Force (FACH), who served as Chief of the Union Glacier Station during the Joint Scientific Expedition (ECA 55) from which soil samples were obtained for the isolation of the micro-organisms used in this study. Sadly, he died in the crash of the Lockheed C-130 Hercules aircraft in the Drake Passage on 9 December 2019) Basonym: *Rhodococcus navarretei* Carrasco *et al.* 2024. The description of this taxon is as given by Carrasco *et al.* [58]. The type strain has a G+C content of 64.5% and genome with a size of ≈5.3 Mbp. Type strain: EXRC-4A-4^T^ (= LMG 33621^T^ = RGM 3539^T^).

### *Hoyosella levis* comb. nov

(le’vis. L. masc. adj. levis, light) Basonym: *Lolliginicoccus levis* Miyanishi *et al.* 2023 The description of this taxon is as given by Miyanishi *et al.* [34]. The type strain has a G+C content of 68% and genome with a size of ≈3.6 Mbp. Type strain: Y7R2^T^ (= KCTC 49749^T^ = NBRC 114883^T^).

## Abbreviations

AAI: average amino acids identity
AF: aligned fraction of orthologous genes
ANI: average nucleotide identity
DDBJ: DNA Data Bank of Japan
EMBL-EBI: European Molecular Biology Laboratory-European Bioinformatics Institute
ENA: European Nucleotide Archive
GTR: General Time-Reversible with empirical base frequencies
ICNP: International Code of Nomenclature of Prokaryotes
ICSP: International Committee on Systematics of Prokaryotes
INSDC: International Nucleotide Sequence Database Collaboration
LPSN: Lists of Prokaryotic names with Standing in Nomenclature (https://www.bacterio.net
Mbp: mega base pairs
ML: maximum likelihood
MLD: maximum likelihood distance
NCBI: National Center for Biotechnology Information
PAM: “Patogénesis Microbiana” isolate collection of Vazquez-Boland’s laboratory
POCP: percentage of conserved proteins
RED: relative evolutionary distance
TaxID: taxonomy identified
WGS: whole genome sequencing
+ASC: ascertainment bias correction
+F: empirical base frequencies
+R: FreeRate model categories.

## Acknowledgements

We thank the University of Edinburgh Digital Research Services and Andy Law for access to, and technical support with Eddie high-performance Linux computing cluster. We also gratefully acknowledge the valuable comments of the expert colleagues with whom we consulted and discussed the concept of using the subgenus category to address the problems associated with the growing genus over-splitting trend.

