## Supplemental data for "Leveraging the subgenus category to address genus over-splitting exemplified with *Prescottella* and other recently proposed *Mycobacteriales* genera"

### SUPPLEMENTAL MATERIAL

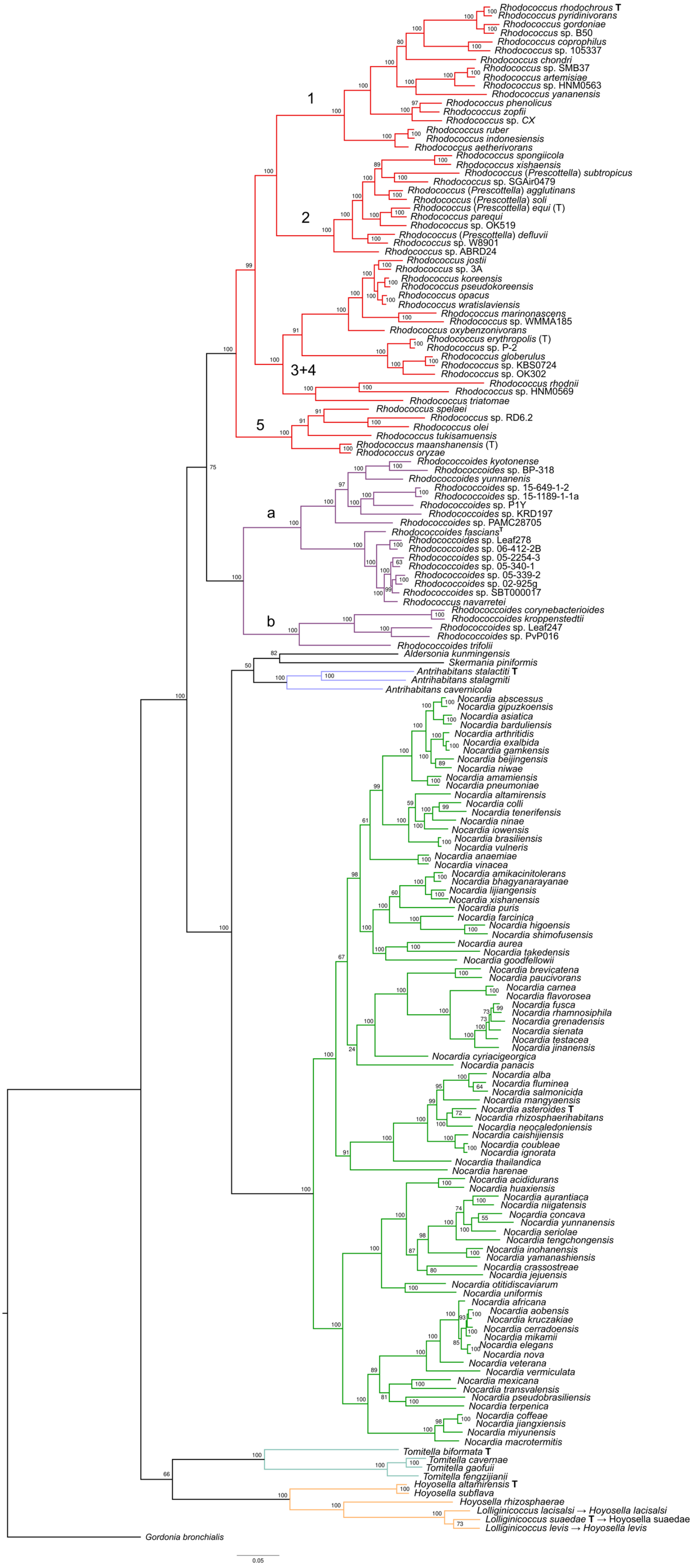


**Fig. S1**

**Fig. S1 (cont.).** Non-normalized tree of the ML phylogeny in **Fig. 1**. Based on a concatenated alignment of 65 conserved proteins selected using the Get_Homologues and Get_Phylomarkers pipelines [1, 2] from 175 *Nocardiaceae*, *Tomitellaceae* and *Hoyosellaceae* species plus *Gordonia bronchialis* DSM 43247$^{T}$ as an outgroup. Accessions for genomes used can be found in Suppl. Dataset, sheet 1. The monotypic, single-strain taxons *Millisia brevis* NBRC 105863$^{T}$ and *Smaragdicoccus niigatensis* DSM 44881$^{T}$ were excluded from the phylogeny because of their uncertain position within the *Nocardiaceae* radiation, which reduced the robustness of the phylogenetic trees (long-branch attraction effect, see ref. [3]). The tree was built using Iqtree v2.0.7 [4] and the substitution model LG+F+R7. Branch colour coding and rhodoccal sugenus/sublineage labels as in **Fig. 1**. The type species of each genus are indicated by a bold-case T, for the proposed *Rhodococcus* subgenera by a T in parenthesis. UltraFastBootstraps values are shown (10,000 replicates). Scale bar, amino acid substitutions per site.Tree plotted using FigTree v1.4.4 (http://tree.bio.ed.ac.uk/software/figtree/).

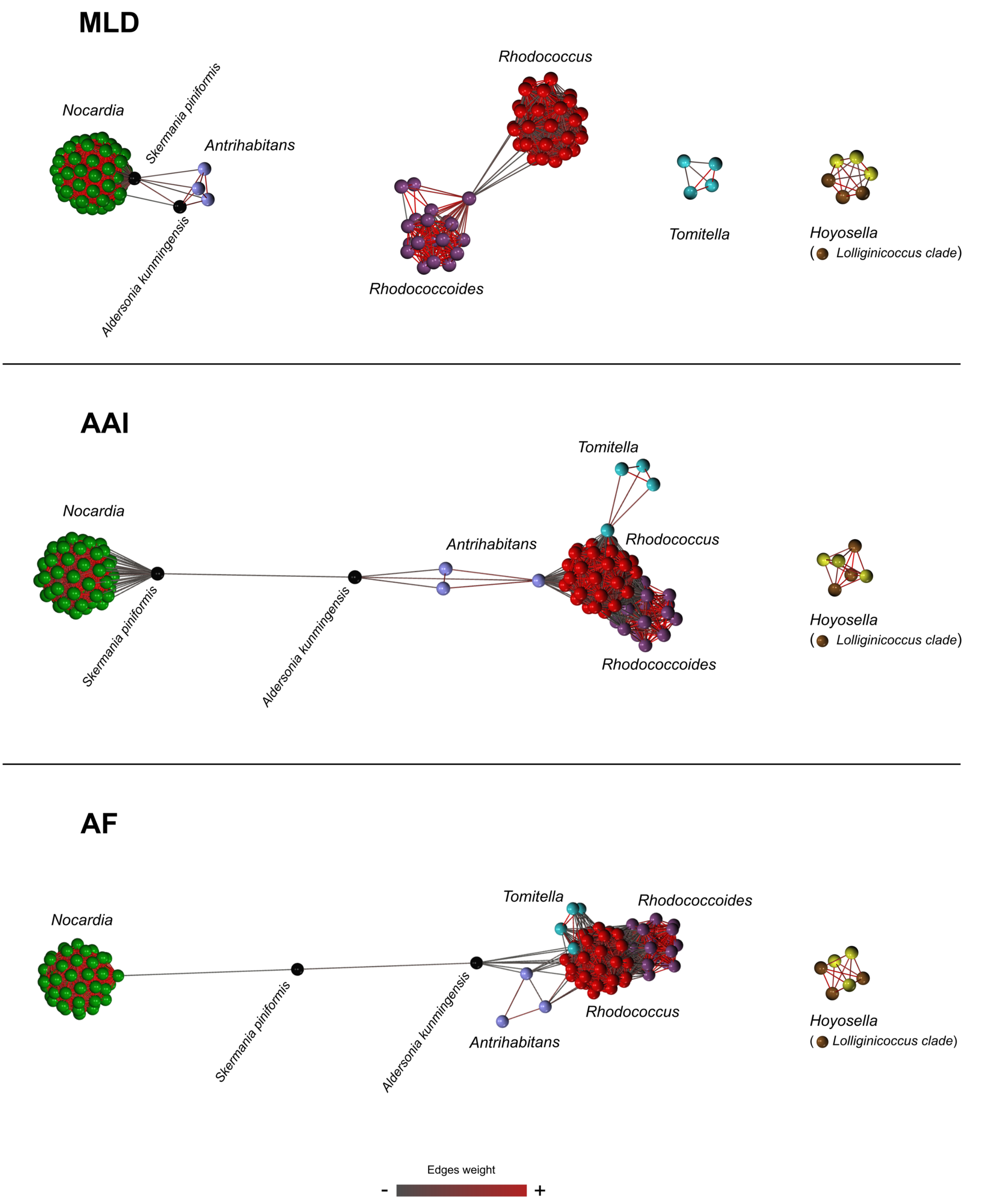


**Fig. S2.** Supragenus-level partitions of **Fig. 2** taxonomic network based on ML distance (MLD) and genome relatedness index (AAI and AF) pairwise comparison matrices. At the applied ct values (MLD = 0.575, AAI = 0.300, AF = 330), all three networks consistently isolate *Hoyosella* (*Hoyosellaceae* family) as an independent cluster. However, *Tomitella* (*Tomitellaceae* family) remains connected with the *Rhodococcus* subnetwork in the AAI and AF graphs (via the type species *Tomitella biformata*). The same happens with the *Antrihabitans* (Nocardiaceae family)

subnetwork. The latter connects with the *Nocardia* subnetwork through *Aldersonia kunmingensis* DSM 45001$^{T}$ *and Skermania piniformis* DSM 43998$^{T}$ which act as bridging nodes denoting their intermediate position between the nocardiae and the rhodococci. Scale shows edge weight from weaker (dark grey) to stronger (red).

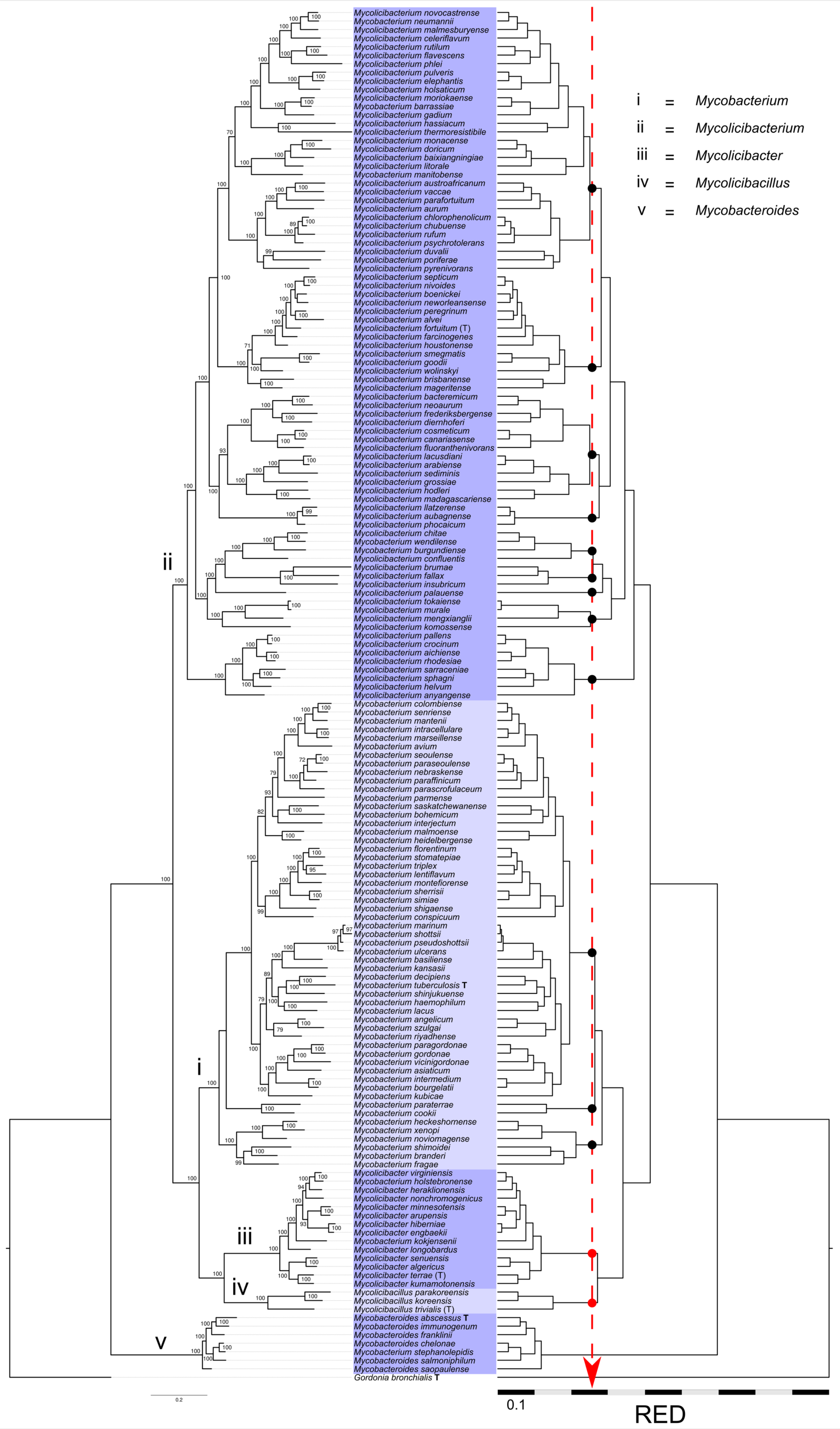
i = Mycobacterium
ii = Mycolicibacterium
iii = Mycolicibacter
iv = Mycolicibacillus
v = Mycobacteroides
Mycolicibacterium novocastrense
Mycobacterium neumannii
Mycolicibacterium malmesburyense
Mycolicibacterium celeriflavum
Mycolicibacterium rutilum
Mycolicibacterium flavescens
Mycolicibacterium phlei
Mycolicibacterium pulveris
Mycolicibacterium elephantis
Mycolicibacterium holsaticum
Mycolicibacterium moriokaense
Mycobacterium barrassiae
Mycolicibacterium gadium
Mycolicibacterium hassiacum
Mycolicibacterium thermoresistibile
Mycolicibacterium monacense
Mycolicibacterium doricum
Mycolicibacterium baixiangningiae
Mycolicibacterium litorale
Mycobacterium manitobense
Mycolicibacterium austroafricanum
Mycolicibacterium vaccae
Mycolicibacterium parafortuitum
Mycolicibacterium aurum
Mycolicibacterium chlorophenolicum
Mycolicibacterium chubuense
Mycolicibacterium rufum
Mycolicibacterium psychrotolerans
Mycolicibacterium duvalii
Mycolicibacterium poriferae
Mycolicibacterium pyrenivorans
Mycolicibacterium septicum
Mycolicibacterium nivoides
Mycolicibacterium boenickei
Mycolicibacterium neworleansense
Mycolicibacterium peregrinum
Mycolicibacterium alvei
Mycolicibacterium fortuitum (T)
Mycolicibacterium farcinogenes
Mycolicibacterium houstonense
Mycolicibacterium smegmatis
Mycolicibacterium goodii
Mycolicibacterium wolinskyi
Mycolicibacterium brisbanense
Mycolicibacterium mageritense
Mycolicibacterium bacteremicum
Mycolicibacterium neoaurum
Mycolicibacterium frederiksbergense
Mycolicibacterium diernhoferi
Mycolicibacterium cosmeticum
Mycolicibacterium canariasense
Mycolicibacterium fluoranthenivorans
Mycolicibacterium lacusdiani
Mycolicibacterium arabiense
Mycolicibacterium sediminis
Mycolicibacterium grossiae
Mycolicibacterium hodleri
Mycolicibacterium madagascariense
Mycolicibacterium llatzerense
Mycolicibacterium aubagnense
Mycolicibacterium phocaicum
Mycolicibacterium chitae
Mycobacterium wendilense
Mycobacterium burgundiense
Mycolicibacterium confluentis
Mycolicibacterium brumae
Mycolicibacterium fallax
Mycobacterium insubricum
Mycolicibacterium palauense
Mycolicibacterium tokaiense
Mycolicibacterium murale
Mycolicibacterium mengxianglii
Mycolicibacterium komossense
Mycolicibacterium pallens
Mycolicibacterium crocinum
Mycolicibacterium aichiense
Mycolicibacterium rhodesiae
Mycolicibacterium sarraceniae
Mycolicibacterium sphagni
Mycolicibacterium helvum
Mycolicibacterium anyangense
Mycobacterium colombiense
Mycobacterium senriense
Mycobacterium mantenii
Mycobacterium intracellulare
Mycobacterium marseillense
Mycobacterium avium
Mycobacterium seoulense
Mycobacterium paraseoulense
Mycobacterium nebraskense
Mycobacterium paraffinicum
Mycobacterium parascrofulaceum
Mycobacterium parmense
Mycobacterium saskatchewanense
Mycobacterium bohemicum
Mycobacterium interjectum
Mycobacterium malmoense
Mycobacterium heidelbergense
Mycobacterium florentinum
Mycobacterium stomatepiae
Mycobacterium triplex
Mycobacterium lentiflavum
Mycobacterium montefiorense
Mycobacterium sherrisii
Mycobacterium simiae
Mycobacterium shigaense
Mycobacterium conspicuum
Mycobacterium marinum
Mycobacterium shottsii
Mycobacterium pseudoshottsii
Mycobacterium ulcerans
Mycobacterium basiliense
Mycobacterium kansasii
Mycobacterium decipiens
Mycobacterium tuberculosis T
Mycobacterium shinjukuense
Mycobacterium haemophilum
Mycobacterium lacus
Mycobacterium angelicum
Mycobacterium szulgai
Mycobacterium riyadhense
Mycobacterium paragordonae
Mycobacterium gordonae
Mycobacterium vicinigordonae
Mycobacterium asiaticum
Mycobacterium intermedium
Mycobacterium bourgelatii
Mycobacterium kubicae
Mycobacterium paraterrae
Mycobacterium cookii
Mycobacterium heckeshornense
Mycobacterium xenopi
Mycobacterium noviomagense
Mycobacterium shimoidei
Mycobacterium branderi
Mycobacterium fragae
Mycolicibacter virginiensis
Mycobacterium holstebronense
Mycolicibacter heraklionensis
Mycolicibacter nonchromogenicus
Mycolicibacter minnesotensis
Mycolicibacter arupensis
Mycolicibacter hiberniae
Mycolicibacter engbaekii
Mycobacterium kokjensenii
Mycolicibacter longobardus
Mycolicibacter senuensis
Mycolicibacter algericus
Mycolicibacter terrae (T)
Mycolicibacter kumamotonensis
Mycolicibacillus parakoreensis
Mycolicibacillus koreensis
Mycolicibacillus trivialis (T)
Mycobacteroides abscessus T
Mycobacteroides immunogenum
Mycobacteroides franklinii
Mycobacteroides chelonae
Mycobacterium stephanolepidis
Mycobacteroides salmoniphilum
Mycobacteroides saopaulense
Gordonia bronchialis T
0.2
0.1
RED

Fig. S3.

**Fig. S3 (cont.)**. ML whole-genome mycobacterial phylogeny. Left, non-normalized tree; right, relative evolutionary distance (RED)-normalized tree. The tree includes 160 mycobacterial species plus *Gordonia bronchialis* DSM 43247$^T$ as an outgroup (see Suppl. Dataset, sheet 1). Phylogeny inferred from a concatenated alignment of universal protein markers using PhyloPhlAn [5]. Tree built with Iqtree v2.0.7 [4] and the substitution model LG+F+R8. The ML tree is fully consistent with previously published mycobacterial phylogenies [3, 6, 7]. The species are named according to NCBI Taxonomy browser based on the five-genus nomenclature proposed by Gupta *et al*. 2018 [6]; the circumscriptions of the five genera are indicated by alternate dark and light blue shade. The RED-normalized tree was partitioned using a taxonomic context-uniform TreeCluster [8] threshold that allowed the separation of the more distal *Mycolicibacter* and *Mycolicibacillus* taxa as independent clusters (red vertical dashed arrow). Application of this clustering cutoff results in significant fragmentation of the other genera proposed by Gupta *et al*. [6], indicating that these authors did not use a consistent, homogeneous taxon demarcation approach; see text for details. Scale bar, amino acid substitutions per site. UltraFastBootstraps values are shown (1,000 replicates). Tree plotted using FigTree v1.4.4 (http://tree.bio.ed.ac.uk/software/figtree/).

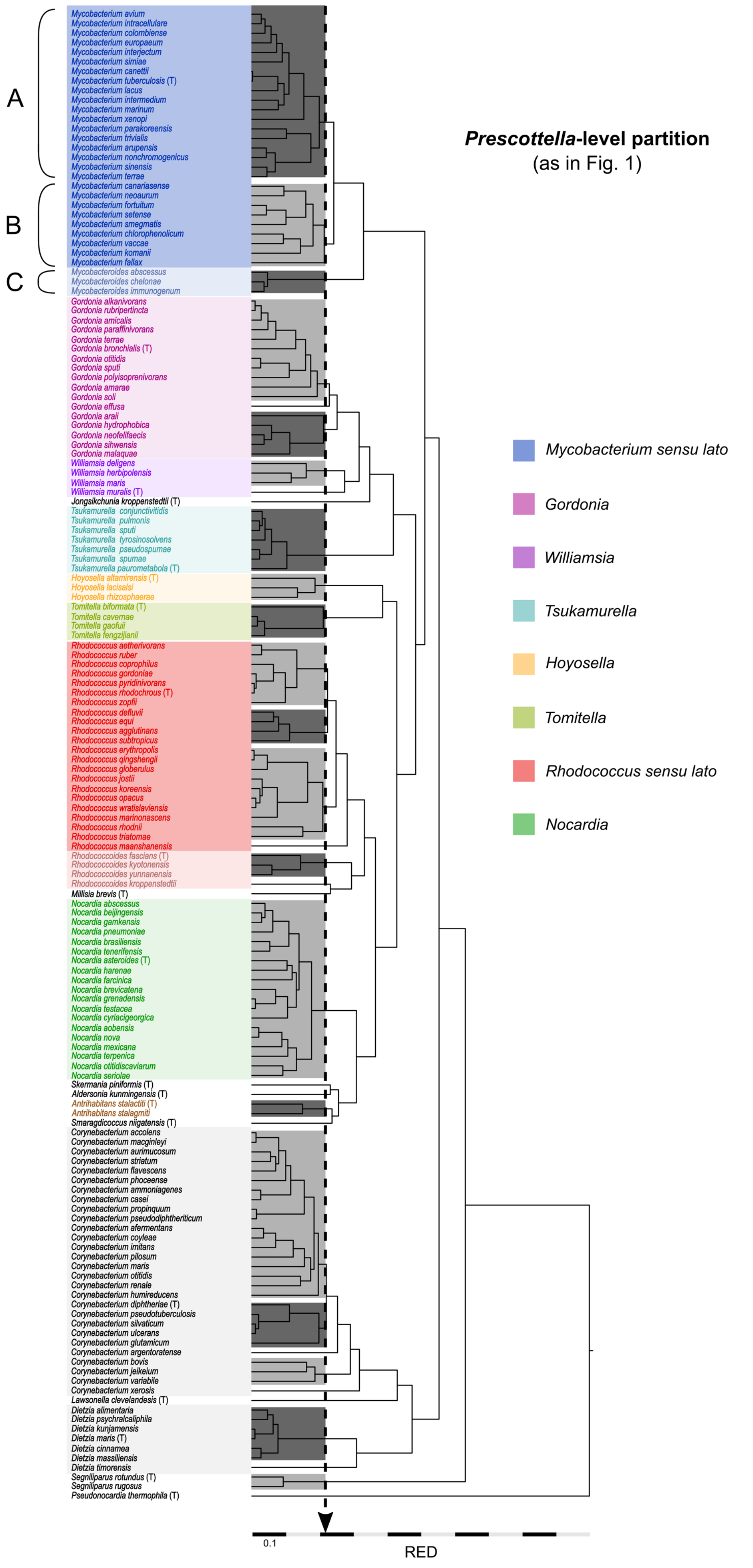

A
B
C
Mycobacterium avium
Mycobacterium intracellulare
Mycobacterium colombiense
Mycobacterium europaeum
Mycobacterium interjectum
Mycobacterium simiae
Mycobacterium canettii
Mycobacterium tuberculosis (T)
Mycobacterium lacus
Mycobacterium intermedium
Mycobacterium marinum
Mycobacterium xenopi
Mycobacterium parakoreensis
Mycobacterium trivialis
Mycobacterium arupensis
Mycobacterium nonchromogenicus
Mycobacterium sinensis
Mycobacterium terrae
Mycobacterium canariasense
Mycobacterium neoaurum
Mycobacterium fortuitum
Mycobacterium setense
Mycobacterium smegmatis
Mycobacterium chlorophenolicum
Mycobacterium vaccae
Mycobacterium komanii
Mycobacterium fallax
Mycobacteroides abscessus
Mycobacteroides chelonae
Mycobacteroides immunogenum
Gordonia alkanivorans
Gordonia rubripertincta
Gordonia amicalis
Gordonia paraffinivorans
Gordonia terrae
Gordonia bronchialis (T)
Gordonia otitidis
Gordonia sputi
Gordonia polyisoprenivorans
Gordonia amarae
Gordonia soli
Gordonia effusa
Gordonia araii
Gordonia hydrophobica
Gordonia neofelifaecis
Gordonia sihwensis
Gordonia malaquae
Williamsia deligens
Williamsia herbipolensis
Williamsia maris
Williamsia muralis (T)
Jongsikchunia kroppenstedtii (T)
Tsukamurella conjunctivitidis
Tsukamurella pulmonis
Tsukamurella sputi
Tsukamurella tyrosinosolvens
Tsukamurella pseudospumae
Tsukamurella spumae
Tsukamurella paurometabola (T)
Hoyosella altamirensis (T)
Hoyosella lacisalsi
Hoyosella rhizosphaerae
Tomitella biformata (T)
Tomitella cavernae
Tomitella gaofuii
Tomitella fengzijianii
Rhodococcus aetherivorans
Rhodococcus ruber
Rhodococcus coprophilus
Rhodococcus gordoniae
Rhodococcus pyridinivorans
Rhodococcus rhodochrous (T)
Rhodococcus zopfii
Rhodococcus defluvii
Rhodococcus equi
Rhodococcus agglutinans
Rhodococcus subtropicus
Rhodococcus erythropolis
Rhodococcus qingshengii
Rhodococcus globerulus
Rhodococcus jostii
Rhodococcus koreensis
Rhodococcus opacus
Rhodococcus wratislaviensis
Rhodococcus marinonascens
Rhodococcus rhodnii
Rhodococcus triatomae
Rhodococcus maanshanensis
Rhodococcoides fascians (T)
Rhodococcoides kyotonensis
Rhodococcoides yunnanensis
Rhodococcoides kroppenstedtii
Millisia brevis (T)
Nocardia abscessus
Nocardia beijingensis
Nocardia gamkensis
Nocardia pneumoniae
Nocardia brasiliensis
Nocardia tenerifensis
Nocardia asteroides (T)
Nocardia harenae
Nocardia farcinica
Nocardia brevicatena
Nocardia grenadensis
Nocardia testacea
Nocardia cyriacigeorgica
Nocardia aobensis
Nocardia nova
Nocardia mexicana
Nocardia terpenica
Nocardia otitidiscaviarum
Nocardia seriolae
Skermania piniformis (T)
Aldersonia kunmingensis (T)
Antrihabitans stalactiti (T)
Antrihabitans stalagmiti
Smaragdicoccus niigatensis (T)
Corynebacterium accolens
Corynebacterium macginleyi
Corynebacterium aurimucosum
Corynebacterium striatum
Corynebacterium flavescens
Corynebacterium phoceense
Corynebacterium ammoniagenes
Corynebacterium casei
Corynebacterium propinquum
Corynebacterium pseudodiphtheriticum
Corynebacterium afermentans
Corynebacterium coyleae
Corynebacterium imitans
Corynebacterium pilosum
Corynebacterium maris
Corynebacterium otitidis
Corynebacterium renale
Corynebacterium humireducens
Corynebacterium diphtheriae (T)
Corynebacterium pseudotuberculosis
Corynebacterium silvaticum
Corynebacterium ulcerans
Corynebacterium glutamicum
Corynebacterium argentoratense
Corynebacterium bovis
Corynebacterium jeikeium
Corynebacterium variabile
Corynebacterium xerosis
Lawsonella clevelandesis (T)
Dietzia alimentaria
Dietzia psychralcaliphila
Dietzia kunjamensis
Dietzia maris (T)
Dietzia cinnamea
Dietzia massiliensis
Dietzia timorensis
Segniliparus rotundus (T)
Segniliparus rugosus
Pseudonocardia thermophila (T)
Prescottella-level partition
(as in Fig. 1)
Mycobacterium sensu lato
Gordonia
Williamsia
Tsukamurella
Hoyosella
Tomitella
Rhodococcus sensu lato
Nocardia
0.1
RED


**Fig. S4**

**Fig. S4**. Tree clustering of the *Mycobacteriales* using *Prescottella* as taxonomic context-uniform subgenus-level partitioning reference [3]. Based on the (RED)-normalized phylogenomic ML tree originally published in ref. [3]. The applied TreeCluster *t* cutoff isolates the main rhodococcal sublineages/proposed *Rhodococcus* subgenera (vertical dashed arrow). This subdivides the mycobacterial tree into three main clusters which only partially correspond to Gupta *et al.* [6] taxons, as follows: (A) *Mycobacterium* + *Mycolicibacter* + *Mycolicibacillus* (B) *Mycolicibacterium*, and (C) *Mycobacteroides.* Figure modified from **Fig. 2** of ref. [3].

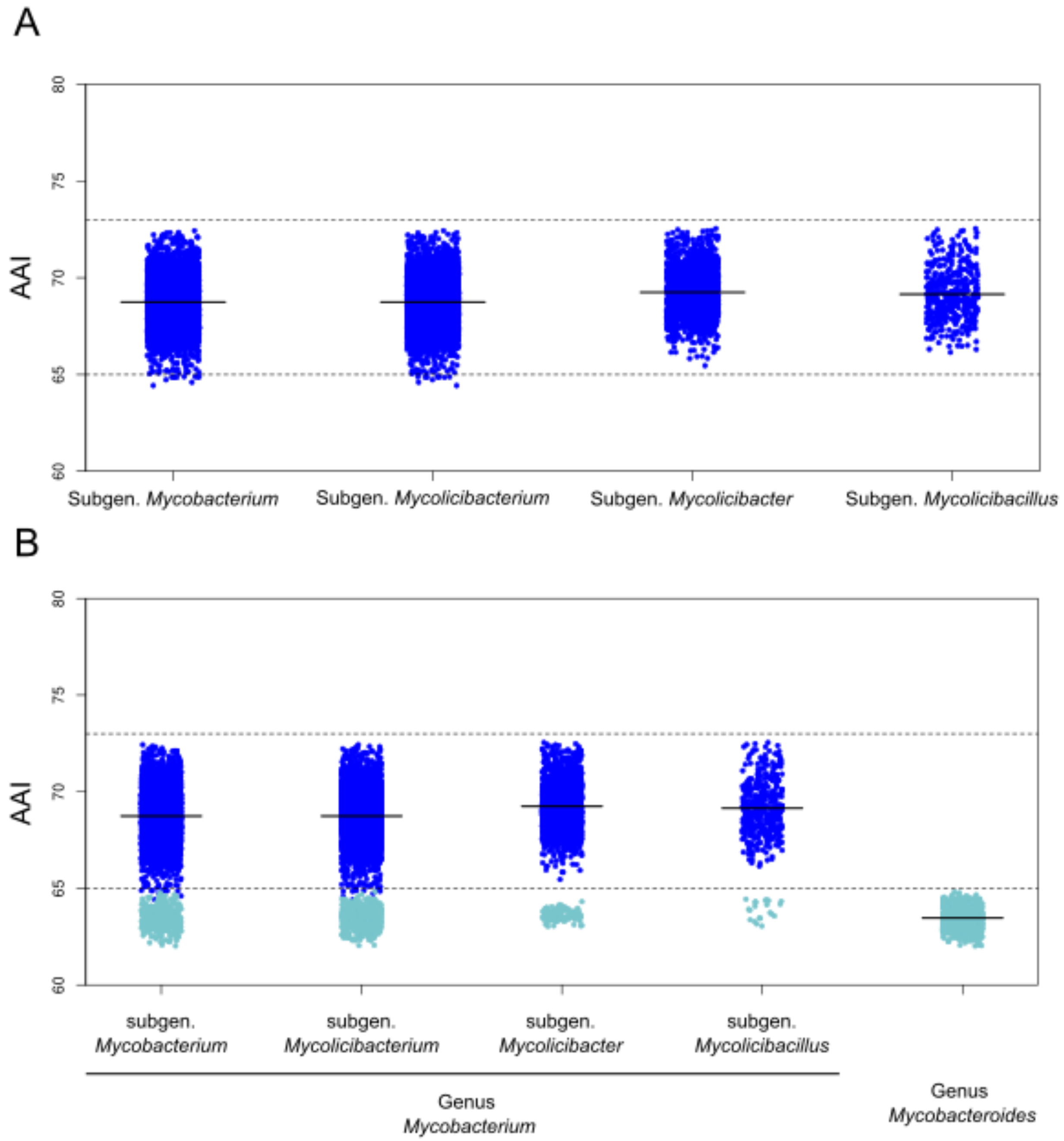


**Fig. S5**. AAI score demarcation of *Mycobacterium* subgenera. The average AAI values are represented by a black horizontal line. (A) scatter plots of *Mycobacterium* AAI inter-subgenus pairwise comparisons. The subgenus demarcation boundary is 73%, the same as the AAI demarcation of rhodoccal subgenera. (B) Same as in A with the AAI comparisons between the species within the four proposed *Mycobacterium* subgenera and the seven species of within the genus *Mycobacteroides* circumscription added (represented as light blue dots). Note the clear-cut separation of the latter below the 65% AAI demarcation standard for genus definition [9-11], supporting the consideration of *Mycobacteroides* as a separate genus rather than a *Mycobacterium* subgenus.
